# Selective Modulator of Nuclear Receptor PPARγ with Reduced Adipogenic Potential Ameliorates Experimental Nephrotic Syndrome

**DOI:** 10.1101/2021.09.14.459479

**Authors:** Claire Burton, Galen Rask, Amanda P Waller, Amy Webb, Marina R Galdino-Pitta, Angelica A. Amato, Rachel Cianciolo, Brian Becknell, Bryce A Kerlin, Francisco A. R. Neves, Alessia Fornoni, Shipra Agrawal

## Abstract

**Background:** Glomerular disease, often manifesting as nephrotic syndrome (NS) with high proteinuria, can be refractory to standard treatment and is typically associated with hypoalbuminemia, hypercholesterolemia and hypercoagulopathy. We hypothesized that the nuclear receptor PPARγ can be selectively modulated using a novel partial agonist, GQ-16, to gain therapeutic advantage over traditional PPARγ agonists (e.g. thiazolidinediones) for the treatment of glomerular disease.

**Methods:** Nephropathy was induced with puromycin amino-nucleoside (PAN) in Wistar rats and treated with Pioglitazone (Pio) or GQ-16. Plasma, serum, and urine chemistries were performed, and kidneys, glomeruli, liver, and white adipose tissue (WAT) were harvested. Lipid accumulation and adipogenic gene expression were measured in adipocytes.

**Results:** PAN-induced proteinuria was significantly reduced with Pio to 64% of PAN-value. It was reduced robustly with GQ-16 to 81% of PAN, which was comparable to controls. While both GQ-16 and Pio restored glomerular *Nphs1* and hepatic *Pcsk9* expression and reduced hypercholesterolemia, GQ-16 also restored glomerular *Nrf2*, and reduced hypoalbuminemia and hypercoagulopathy. Furthermore, RNA-seq analysis identified both common and distinct restored glomerular genes downstream of Pio and GQ-16. Pio but not GQ-16 significantly induced *aP2* (fatty acid binding protein) in adipocytes and in WAT. Pio induced more lipid accumulation than GQ-16 in differentiated adipocytes. Both, Pio and GQ-16 induced insulin sensitizing adipokines in WAT with varying degrees.

**Conclusions:** Selective modulation of PPARγ by a partial agonist, GQ-16, is more advantageous than pioglitazone in reducing proteinuria and NS associated co-morbidities, while reducing the adipogenic side-effects conferred by traditional PPARγ full agonists.

**Translational Statement:** The authors have previously reported that type-II diabetes drugs, thiazolidinediones (PPARγ agonists), also provide beneficial effects in reducing podocyte and glomerular injury. However, these drugs are associated with adverse effects such as weight gain, and their effects on glomerular disease-associated features are largely unexplored. Their current findings demonstrate that PPARγ can be selectively modulated by its partial agonist, GQ-16, which reduces proteinuria and improves nephrotic syndrome (NS) with reduced side-effects typically conferred by thiazolidinediones. These findings not only deepen our molecular understanding of the role of PPARγ in glomerular disease and underscore the potential for partial agonists of PPARγ, such as GQ-16 as a treatment modality for NS, but also lend the possibility of its potential benefits in diabetic nephropathy.

## Introduction

Various forms of glomerular disease, manifesting as nephrotic syndrome (NS) with high-grade proteinuria, can be frequently refractory to treatment leading to progression to chronic kidney disease and end-stage kidney disease (ESKD) ^1–3^. Additionally, NS is typically associated with edema, hypoalbuminemia, hypercholesterolemia, systemic immune dysregulation, and hypercoagulopathy ^4–8^. In order to identify effective treatments for glomerular disease, we and others have previously reported that peroxisome proliferator-activated receptor γ (PPARγ) agonists and thiazolidinediones (TZDs) such as pioglitazone (Pio), directly protect podocytes from injury ^9–12^ and reduce proteinuria and glomerular injury in various animal models of glomerular disease ^13–21^. They have also been shown to improve clinical outcomes in NS patients refractory to steroid treatment ^13, 22^. Moreover, these protective effects in experimental models have been shown to be mediated by activation of podocyte PPARγ, thus indicating a pivotal role for PPARγ in maintaining glomerular function through the preservation of podocytes even in non-diabetic glomerular diseases in addition to their general beneficial metabolic, insulin-sensitizing and anti-inflammatory effects ^13, 14, 20, 21^.

Since the identification of PPARs in 1990, PPARγ has been recognized as a nuclear receptor superfamily member, a ligand-dependent transcription factor, and a master regulator of adipogenesis and metabolism. The ability of PPARγ to regulate adipogenesis and lipid storage accounts for the insulin sensitizing effects of its agonists or anti-diabetic drugs known as TZDs ^23^. Interestingly, in patients with diabetic nephropathy (DN), TZDs have been shown to exhibit antiproteinuric effects in a meta-analyses study as well as a decrease in urinary podocyte loss ^24, 25^. Moreover, PPARγ can exist in tissue-specific and function-specific forms which can be generated due to alternative splicing and promoter usage ^26, 27^ as well as its differential phosphorylation, specifically at Serine (Ser) 273 ^28, 29^, which have been shown to be important determinants of its effects on adipogenesis and insulin sensitivity (Fig. 1). However, the roles of PPARγ alternative splicing or differential phosphorylation in glomerular disease are unexplored.

**Figure 1.**
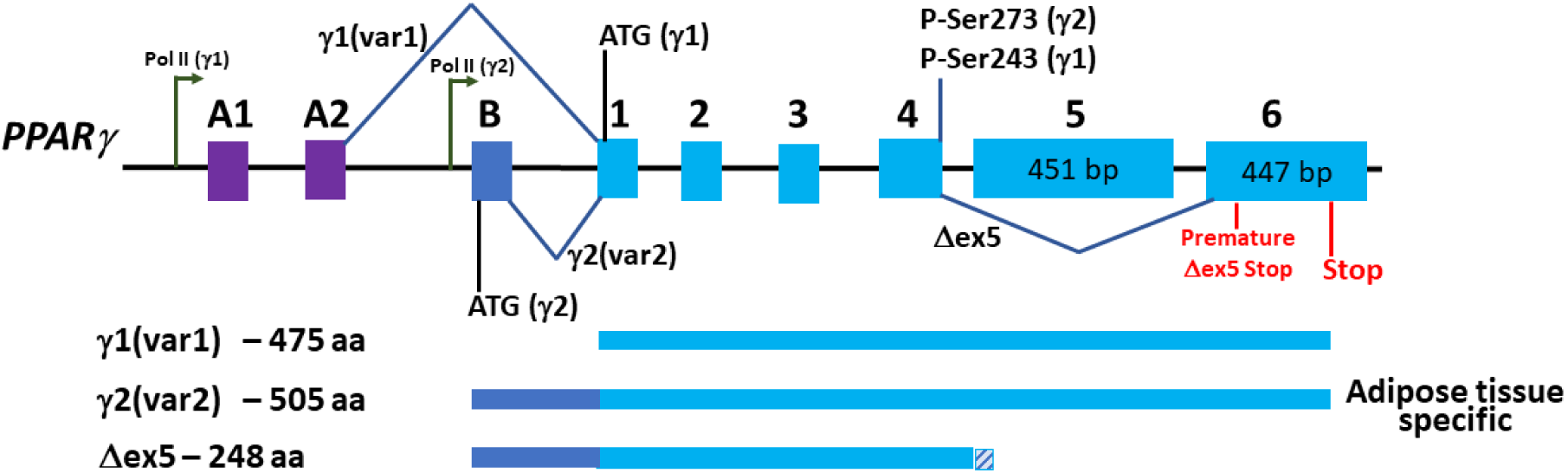
PPARγ Isoforms and Phosphorylation Site. PPARγ exists in mainly two major isoforms, y1 and y2, which are a result of different promoter usage by RNA Polymerase II (depicted in tall arrows) as well as alternative splicing (depicted by exon skipping), resulting in variants 1 and 2. While, y1 variant 1 includes exons A1 and A2 upstream of the start codon (ATG) on exon 1, y2 variant 2 contains the transcription start site preceding exon B and initiates translation on the start codon (ATG) in exon B, resulting in y2 protein product, which is 30 amino acids longer than y1. In a relatively unexplored variant that skips exon 5 (Δex5), a frameshift occurs on exon 6 due to exon 5 being spliced out, which creates a premature stop codon. One of the major sites of phosphorylation on Serine 273 (Ser273) is depicted. This is coded by the last codon on exon 4, thus it remains conserved in variant1, variant 2, and Δex5. The relative position of Ser273 is changed to Ser243 in the isoform γ1. This figure depicts the human annotations for variants 1 (var1: NM_138712) and 2 (var2: NM_015869) encoding isoforms γ1 and γ2. These variants and isoforms correspond to reversed numbers in rat and are annotated as 2 (var2: NM_001145366) and 1 (var1: NM_013124), respectively.

Targeting PPARγ via two widely marketed anti-diabetic drugs, the TZDs Pio and especially rosiglitazone, has been under recent scrutiny because of reported side effects such as weight gain, increased risk of edema, heart failure, bone loss, and bladder cancer ^30–35^. More recently, breakthrough discoveries in the field of PPARγ biology have led to the generation of a series of novel compounds with a weak traditional PPARγ agonistic activity (adipogenic), but very good anti-diabetic activity ^28, 36–38^. GQ-16, a novel selective modulator of PPARγ has been demonstrated to improve insulin sensitivity in diabetic mice in the absence of weight gain and edema ^37, 38^. Moreover, GQ-16 treatment is accompanied by reduced activation of *aP2* and lipid accumulation in vitro and induction of thermogenesis-related genes in epididymal fat depots in vivo, suggesting that browning of visceral WAT may have contributed to weight loss ^37, 38^. This advantageous pharmacological profile appears to be due to the unique binding mode of GQ-16 to PPARγ and stabilization of beta sheets, which is distinct from traditional TZDs ^37, 38^.

Based on the above, we hypothesized that the downstream effects of PPARγ can be mechanistically dissociated and that selected manipulation of PPARγ by a novel partial agonist, GQ-16, will result in a better and targeted therapeutic advantage in glomerular disease. To test this hypothesis, we analyzed the abilities of GQ-16 and Pio to: 1) provide reduction in proteinuria and glomerular injury, 2) regulate systemic and adipogenic effects by complete vs. partial agonism of PPARγ, and 3) modulate downstream molecular pathways, in an animal model of glomerular disease.

## Methods

### Animal Studies: Glomerular Disease Model and Treatment with GQ-16 and Pio

This study was approved by the Institution Animal Care and Use Committee at Nationwide Children’s Hospital, and their guidelines were followed when performing the experiments. GQ-16 was synthesized as previously described and quality tested for 99% purity ^37^. Male Wistar rats were intravenously (IV) injected with puromycin aminonucleoside (PAN) (Sigma-Aldrich, St. Louis, MO) and treatment groups were treated by oral gavage with Pioglitazone (Alfa Aesar, Tewksbury, MA), GQ-16, or a sham vehicle daily. GQ-16 and Pio dosages were determined based on *in vitro* PPARγ activation, *aP2* induction and lipid accumulation data in the current study and our previous studies on the expression of thermogenesis-related genes by GQ-16 ^38^ and proteinuria reducing beneficial effects by Pio ^13^. Serum and urine chemistries were performed, and kidneys, glomeruli, liver, and white adipose tissue (WAT) were harvested for histology, RNA, and protein extraction. Blood was collected through the inferior vena cava, then processed to Platelet Poor Plasma (PPP) as described previously for determination of thrombin generation parameters ^39, 40^. Detailed methods are provided in Supplementary.

### Urinalysis and Serum Chemistry

Urine was analyzed on gel and protein:creatinine ratio (UPCr) analyses were performed by Antech Diagnostics GLP (Morrisville, NC) ^13, 39^. Serum albumin and cholesterol were measured at the Clinical Pathology Services, The Ohio State University’s College of Veterinary Medicine.

### Coagulopathy Measurement

Thrombin generation assays (TGA) were performed on PPP samples collected from the rats to determine various parameters such as the endogenous thrombin potential (ETP) and peak thrombin concentration.

### RNA Isolation, Quantitative Real Time Reverse Transcription-Polymerase Chain Reaction

Total RNA was extracted from isolated glomeruli, liver, and WAT epididymal fat tissue and subjected to qRT-PCR using gene specific primers (Table 1).

**Table 1.**
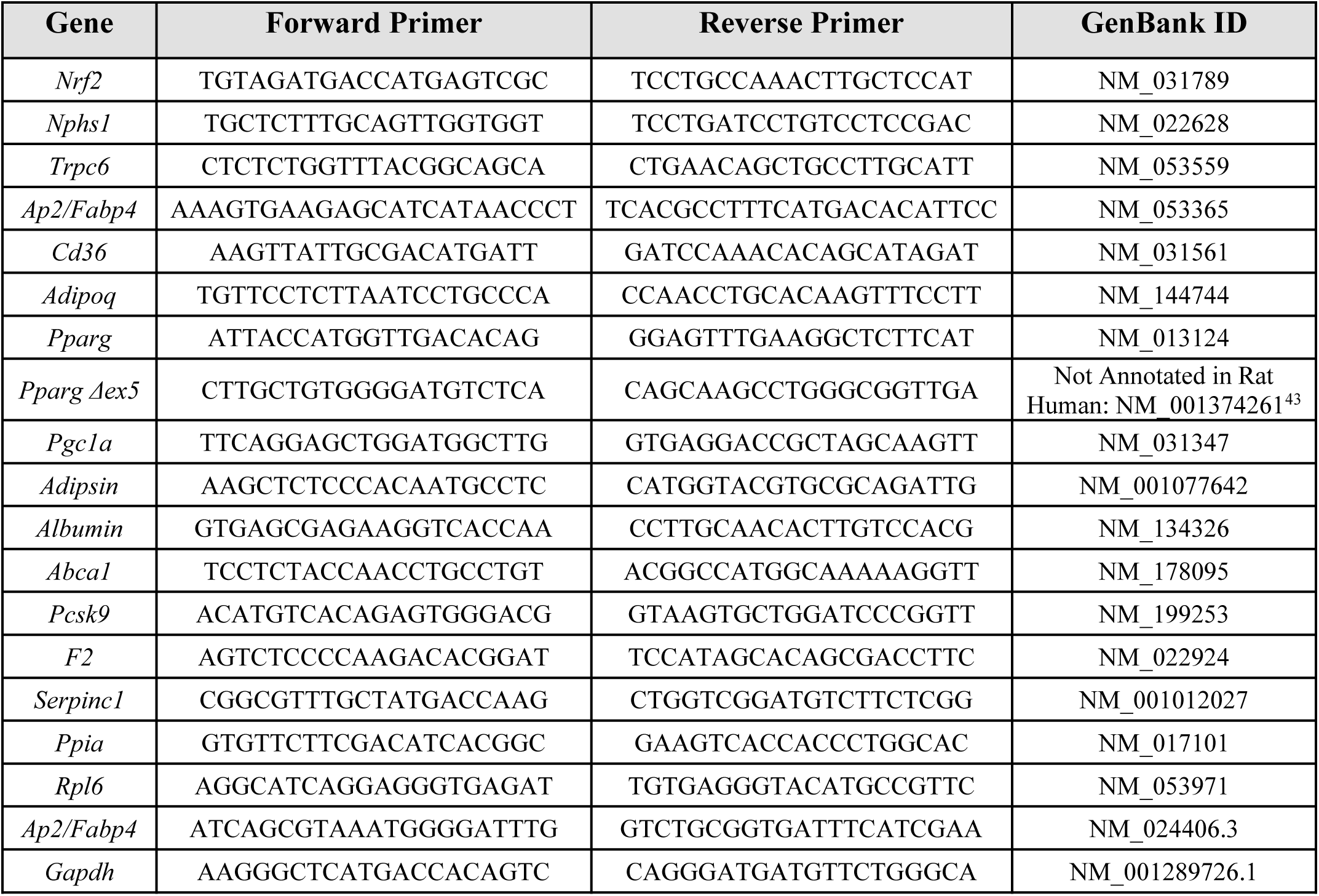
Genes and Their Respective Primers Used in This Study (5’ to 3’ direction)

### RNA Sequencing, Differential Gene Expression, Pathway Analysis and Ontology Enrichment

Glomerular RNA was subjected to RNA sequencing using NovaSeq6000 SP followed by analysis of differentially expressed genes (DEGs), venn diagram and heat map generation, pathway analysis and ontology enrichment.

### Protein Isolation, SDS-PAGE and Western Blotting

Protein was isolated from glomeruli, liver, and WAT epididymal fat tissues and characterized on western blot using primary antibodies for anti-Phospho-PPARγ-Ser 273 (Bioss, Woburn, MA), anti-PPARγ (Proteintech, Carlsbad, CA), and anti-GAPDH (Cell Signaling, Danvers, MA). Densitometry was performed on the bands using ImageJ (National Institutes of Health, Bethesda, MD) software.

### PPARγ Activation Luciferase Assay

HeLa cells were transiently co-transfected with plasmids containing human *PPARG* Ligand-Binding Domain fused to the *GAL4* DNA-Binding Domain and a plasmid containing the luciferase reporter gene under the regulation of five Gal4 DNA-binding elements, treated with varying concentrations of Pio or GQ-16, and assayed for luciferase activity.

### 3T3-L1 Preadipocyte Culture and Differentiation, Lipid Accumulation Assessment and Ap2 Induction

3T3-L1 preadipocytes were cultured and differentiated, treated with varying concentrations of Pio and GQ-16 and assayed for gene expression analysis and intracellular lipid accumulation.

### PPAR-Responsive Element Prediction

A PPARgene database (ppargene.org) was used to predict the sequence specific PPAR-responsive elements (PPRE) on all the target genes measured in this study ^41^.

### Statistical Analysis

Statistics were performed using the GraphPad Prism software version 8.2.0 for Windows (GraphPad Software, San Diego, CA). Data were expressed as mean ± standard error of mean and compared using analysis of variance (ANOVA) followed by the Tukey post-hoc for grouped comparisons and Mann-Whitney test or Student’s *t*-test for pairwise comparisons. Linear regression was used to quantify the correlation of measurement values using the GraphPad Prism software version 8.2.0 for Windows (GraphPad Software, San Diego, CA). *P* value significance was depicted as: * *p* <0.05, ** *p* <0.01, *** *p* <0.001, **** *p* <0.0001.

## Results

### GQ-16 Activates PPARγ Partially and Reduces Proteinuria and Hypoalbuminemia in a Rat Model of PAN-Induced Nephropathy

In order to assess the efficacy of GQ-16 and to compare it with a traditional agonist of PPARγ, Pio, in reducing proteinuria in a glomerular disease model, a PAN-induced nephropathy model was utilized. Efficacy of PPARγ activation was first measured and compared *in vitro* using a luciferase assay system. In agreement with previous data, GQ-16 displayed partial PPARγ agonist activity in transactivation assays (Fig. S1)^37^. It activated PPARγ in a dose-dependent manner and elicited ∼ 30-50% of the maximal activity induced by the full agonist Pio (Supplementary Fig. S1). A single IV PAN injection of 50 mg/kg to male Wistar rats induced massive proteinuria on Day 11 (32.5 ± 9.3 mg/mg; *p* = 0.0006), which started appearing on Day 4 (Fig. 2A and B). Control rats, which received IV saline injection, maintained baseline levels of urinary protein (2.3 ± 0.2 mg/mg). Daily Pio treatment resulted in a significant mean reduction in PAN-induced proteinuria to 64% (13.2 ± 5.5 mg/mg; *p* = 0.05). Interestingly, GQ-16 treatment decreased PAN-induced proteinuria more robustly to 81% reduction (8 ± 3.5 mg/mg; *p* = 0.004) (Fig. 2A and B). Additionally, the proteinuria levels with GQ-16 treatment were comparable to control levels, while Pio treatment remained significantly different from Control (*p* = 0.004).

**Figure 2.**
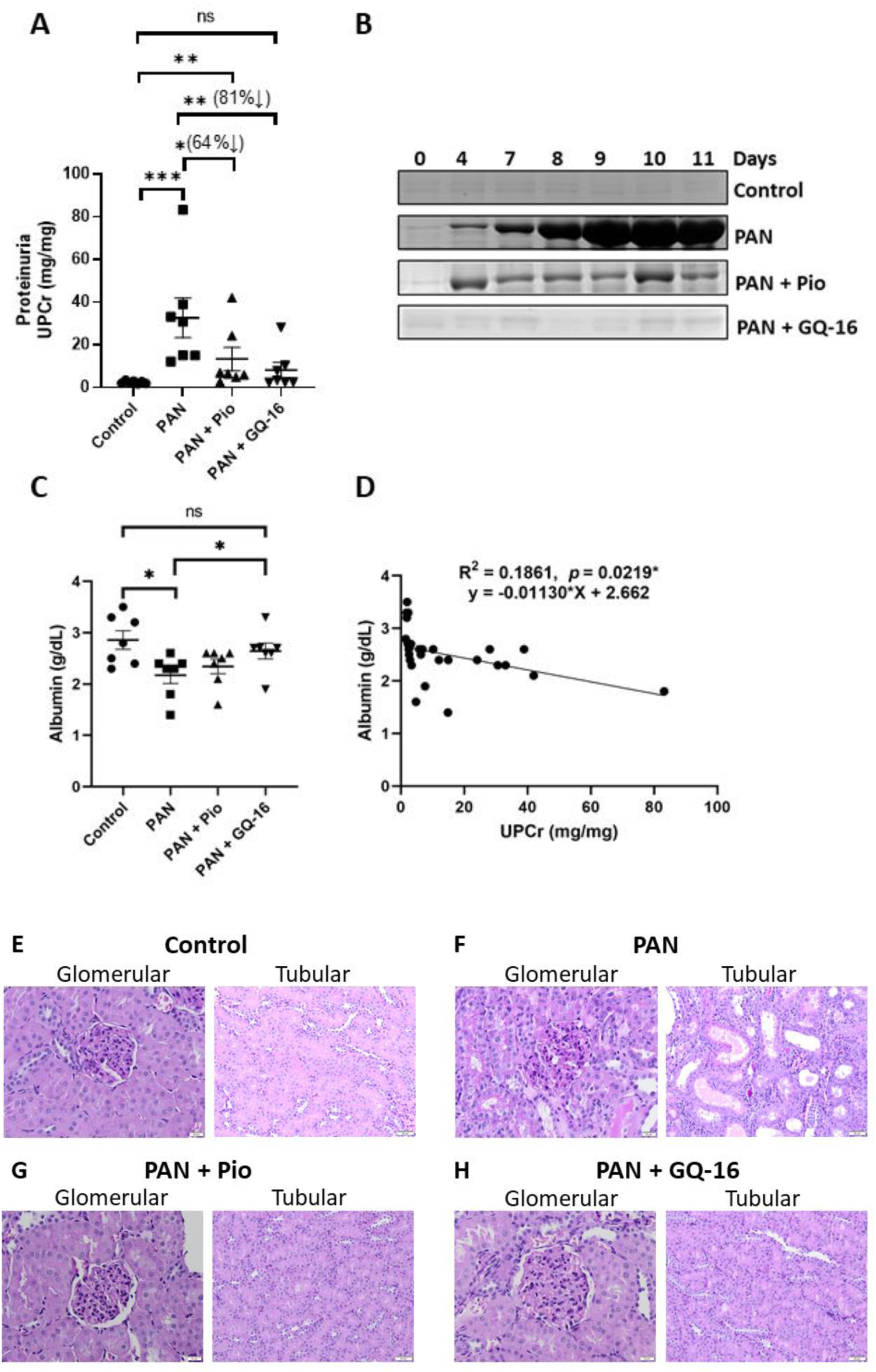
GQ-16 Reduces Proteinuria and Hypoalbuminemia in a Rat Model of PAN-Induced Nephropathy. Proteinuria was induced in male Wistar rats after single intravenous injection of PAN (50 mg/kg) on Day 0. Treatment groups received pioglitazone (Pio, 10mg/kg; n=7) or GQ-16 (40mg/kg; n=7) daily by oral gavage for 11 days. **(A)** Urinary protein/creatinine ratios (UPCr) were plotted from Day 11 urine samples (\**p* <0.05, ** *p* <0.01, *** *p* <0.001, ns= *p* >0.05). **(B)** Representative gels showing urinary albumin bands after PAN injections and treatments with Pio and GQ-16. Equal volumes (5 µl) of urine from selected days were analyzed by SDS-polyacrylamide gel electrophoresis and Coomassie Blue staining. **(C)** Serum albumin concentrations from Day 11 were plotted for the four groups (\**p* <0.05, ns = *p* >0.05). **(D)** Linear regression was performed on all the rats combined which showed correlation of serum albumin with proteinuria. **(E-H)** Histologic evaluation of kidneys stained with periodic acid-Schiff method. PAN-injected rats (**F**) had dilated tubules that contained protein casts. These were not present in saline-injected rats (**E**) or in PAN injected rats that were treated with Pio (**G**) or GQ-16 (**H**). As expected in the model of minimal change disease, the glomeruli were normal.

Assessment of serum albumin levels in these rats showed a decrease in PAN injected rats as compared to control rats (2.2 ± 0.2 g/dL vs. 2.9 ± 0.2 g/dL; *p* = 0.02). Treatment with daily Pio resulted in a modest but insignificant increase in serum albumin (2.3 ± 0.1 g/dL; *p* = 0.26), while treatment with GQ-16 daily resulted in a significant increase in serum albumin levels (2.6 ± 0.1 g/dL; *p* = 0.014), which were comparable to control (*p* = 0.63, ns) (Fig. 2C). Additionally, serum albumin showed a significant correlation to proteinuria in all the rats combined (*p* = 0.02) (Fig. 2D).

Histologic evaluation of kidneys from PAN rats revealed numerous dilated tubules with intratubular protein casts and minimal glomerular lesions as expected with the PAN model (Fig. 2E and F). Some tubules were lined by attenuated epithelium whereas others had hypertrophied epithelial cells, and there was mild multifocal lymphocytic inflammation. Tubular casts, tubular dilation, epithelial cell attenuation and hypertrophy and inflammation were all prevented by both Pio and GQ-16 treatments (Figures 2G and H, respectively

### Glomerular Gene Expression of Podocyte Marker Nphs1 and PPARγ Target Gene Nrf2 is Restored with Treatment with GQ-16

Since podocytopathy is a characteristic of proteinuria in glomerular disease and to understand the role of PPARγ activation in altering glomerular pathways, we measured the expression of relevant podocyte markers and genes in the glomeruli of nephrotic and treated rats. PAN-induced nephropathy resulted in a reduction in the glomerular expression of podocyte marker *Nphs1* (encoding for nephrin), a critical component of slit diaphragm, as well as *Nrf2* (encoding for nuclear factor erythroid 2-related factor 2), a target gene downstream of *PPAR*γ (Fig. 3A-D). *Nphs1* expression was down-regulated with PAN and significantly restored with GQ-16 treatment and modestly with Pio treatment (Fig. 3A). *Nphs1* expression also correlated with reduction in proteinuria in these rats (*p* = 0.01) (Fig. 3B). Notably, while GQ-16 treatment resulted in marked restoration of *Nrf2* expression, Pio treatment did not (Fig. 3C), and *Nrf2* expression levels correlated strongly with reduction in proteinuria (*p* = 0.001) (Fig. 3D). While *Trpc6* (encoding for transient receptor potential cation channel, subfamily C, member 6) showed a trend towards induction with PAN and reduction with both Pio and GQ-16 treatments, its correlation with proteinuria was not found to be significant (Fig. 3E and F). Furthermore, we found that our gene expression data corroborated the prediction of PPREs on the target genes measured in this study using the ‘PPARgene’ database (Table 2) ^41^. This database was developed using a machine learning method to predict novel PPAR target genes by integrating in silico PPRE analysis with high throughput gene expression data. For example, both *Nrf2* and *Nphs1* are predicted to contain PPREs flanking their transcription start sites.

**Figure 3.**
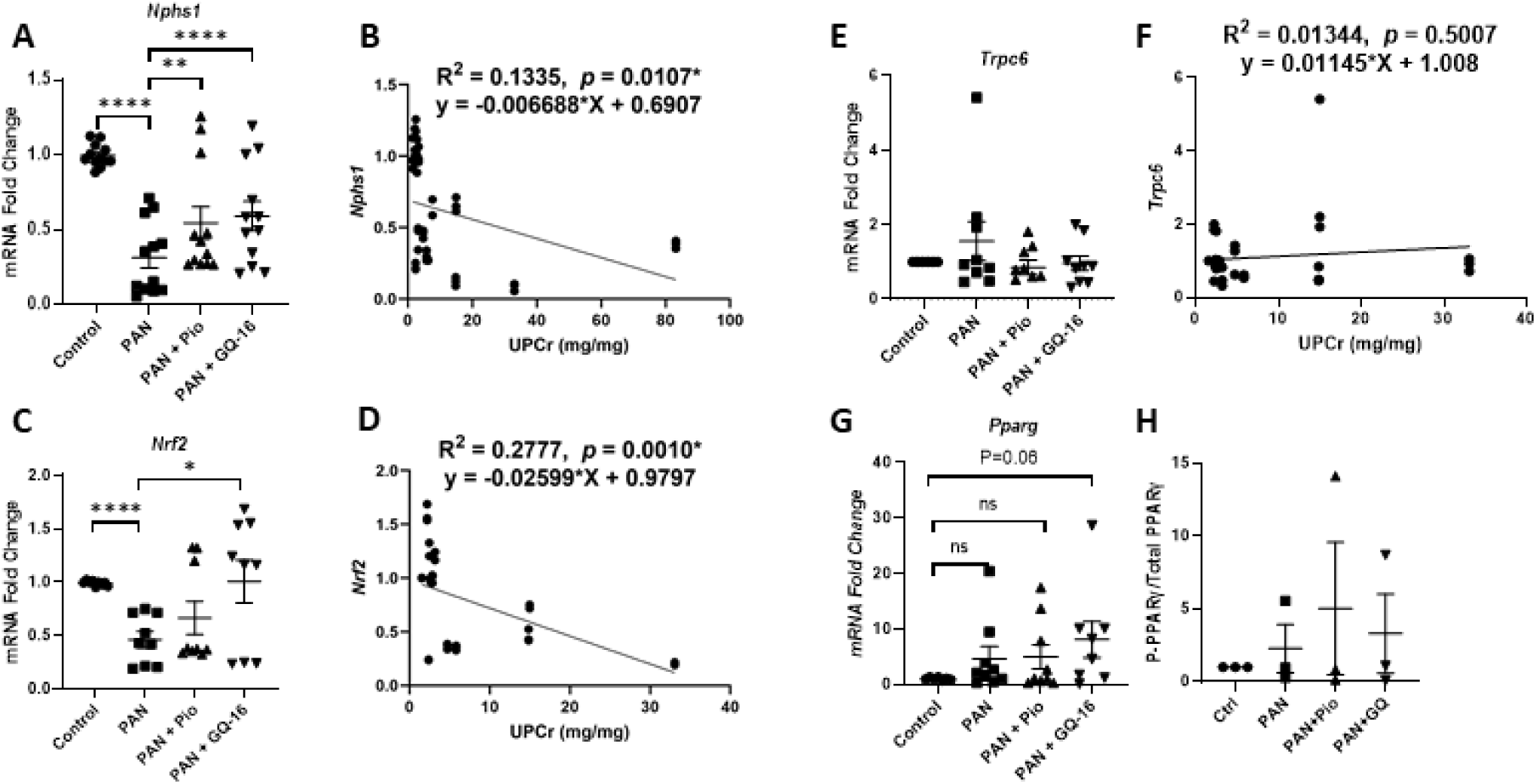
Glomerular Gene Expression of Podocyte Marker *Nphs1* and PPARγ Target Gene *Nrf2* is Restored with Treatment. Glomerular gene expression was measured by real time RT-PCR from total RNA extracted from the glomeruli isolated from Control, PAN-injected, and PAN-injected rats treated with Pio or GQ-16 ((n ≥ 3/group, assay in triplicates). Mean ± SEM plotted of **(A)** *Nphs1,* **(C)** *Nrf2,* **(E)** *Trpc6,* and **(G)** *Pparg.* \**p* <0.05, ** *p* <0.01, *** *p* <0.001, **** *p* <0.0001. Gene expression was normalized to house-keeping gene *Rpl6.* Linear regression analyses were performed to correlate proteinuria with **(B)** *Nphs1,* **(D)** *Nrf2,* and **(F)** *Trpc6* expression. **(H)** Phosphorylated-PPARγ at Serine 273 was measured and normalized to total PPARγ using total protein isolated from Control, PAN-injected, and PAN-injected rats treated with Pio and GQ-16 from glomeruli and plotted.

**Table 2.** PPAR-Responsive Element (PPRE) Prediction in Target Genes.

| Gene | Tissue (Current Study) | *p* Value* | Confidence Level | PPREs Predicted<sup>#</sup> |
| --- | --- | --- | --- | --- |
| *Nrf2* | Glomerular | 0.45663 | low | 7 |
| *Nphs1* | Glomerular | 0.63696 | medium | 4 |
| *Trpc6* | Glomerular | – | – | – |
| *Ap2/Fabp4* | WAT | 0.99997 | high | 10 |
| *Cd36* | WAT | 0.97886 | high | 3 |
| *Adipoq* | WAT | 0.97153 | high | 12 |
| *Pparg* | Glomerular/WAT | 0.91081 | high | 2 |
| *Pgc1a* | WAT | – | – | – |
| *Adipsin* | WAT | 0.96216 | high | 6 |
| *Albumin* | Hepatic | 0.63443 | medium | 2 |
| *Abca1* | Hepatic | 0.99483 | high | 3 |
| *Pcsk9* | Hepatic | – | – | – |
| *F2* | Hepatic | 0.55991 | low | 4 |
| *Serpinc1* | Hepatic | – | – | – |
| *Ppia* | Housekeeping (WAT, hepatic) | – | – | – |
| *Rpl6* | Housekeeping (glomerular) | – | – | – |

\*Probability of being a PPAR target gene, higher value means a higher confidence
High-confidence ( $p > 0.8$ ), median-confidence ( $0.8 \geq p > 0.6$ ), low-confidence category ( $0.6 \geq p > 0.45$ ).
Genes with $p$ value $\leq 0.45$ were predicted as negative.
<sup>#</sup>Putative PPREs in the 5Kb transcription start site (TSS) flanking region

Next, we determined the ability of Pio and GQ-16 to alter *Pparg* gene expression and its phosphorylation status at the Serine 273 position. Phosphorylation of PPARγ at Ser273 has been shown to be an important determinant of its activity for its insulin sensitizing effects ^28, 29^, and thus to understand its role in the reduction of proteinuria, we measured its levels in the glomeruli of nephrotic and Pio and GQ-16-treated rats. While *Pparg* expression tended to be greater in PAN and PAN + Pio groups compared to control, and somewhat increased with GQ-16 treatment, it was not significant (*p* = 0.06) (Fig. 3G) and the overall expression in all the rats did not correlate with proteinuria (data not shown). Furthermore, the phosphorylation status of PPARγ at Ser273 position was unaltered with Pio and GQ-16 in PAN-nephrotic rats (Supplementary Fig. S2 and Fig. 3H), and it did not correlate with proteinuria.

### RNASeq Analysis Reveals that Pio and GQ-16 Restore Common and Distinct Glomerular Genes, Pathways and Downstream Processes

Injury with PAN resulted in 1089 DEGs compared to controls, and 26 of these DEGs were restored by both Pio and GQ-16 treatments, whereas 106 unique GQ-16 regulated DEGs and 17 unique Pio-regulated DEGs were identified (Fig. 4A). Overall, Pio and GQ-16 treatment resulted in 75 and 173 DEGs compared to PAN injury, which included 29 common and 190 distinct genes (Fig. 4A). Of the 29 common DEGs, 28 DEGs were down-regulated by both Pio and GQ-16 and only 1 DEG upregulated by both treatments when compared to PAN (Fig. 4B). Of the 190 distinct DEGs identified between Pio and GQ-16 treatments, 41 were down-regulated and 5 up-regulated by Pio and 124 down-regulated and 20 up-regulated by GQ-16 treatment. Fig. 4C depicts a heatmap of all the DEGs between PAN vs. Control and those restored by (i) only GQ-16, or (ii) both or (iii) only Pio treatments. IPA of DEGs between PAN+Pio vs. PAN (75 DEGs) and PAN+GQ-16 vs. PAN (173 DEGs) revealed the top canonical glomerular pathways that are associated with the proteinuria reducing beneficial effects of Pio and GQ-16, respectively (Figs. 4D and E). These include the IL-8, IL-12 and NFκB pathways for Pio and cell cycle regulation and matrix metalloprotease pathways for GQ-16. Furthermore, ontology enrichment analysis identified the top biological processes (Supplementary Figs. S3A and B), cellular components (Supplementary Figs. S3C and D) and molecular functions (Supplementary Figs. S3E and F) associated with each treatment.

**Figure 4.**
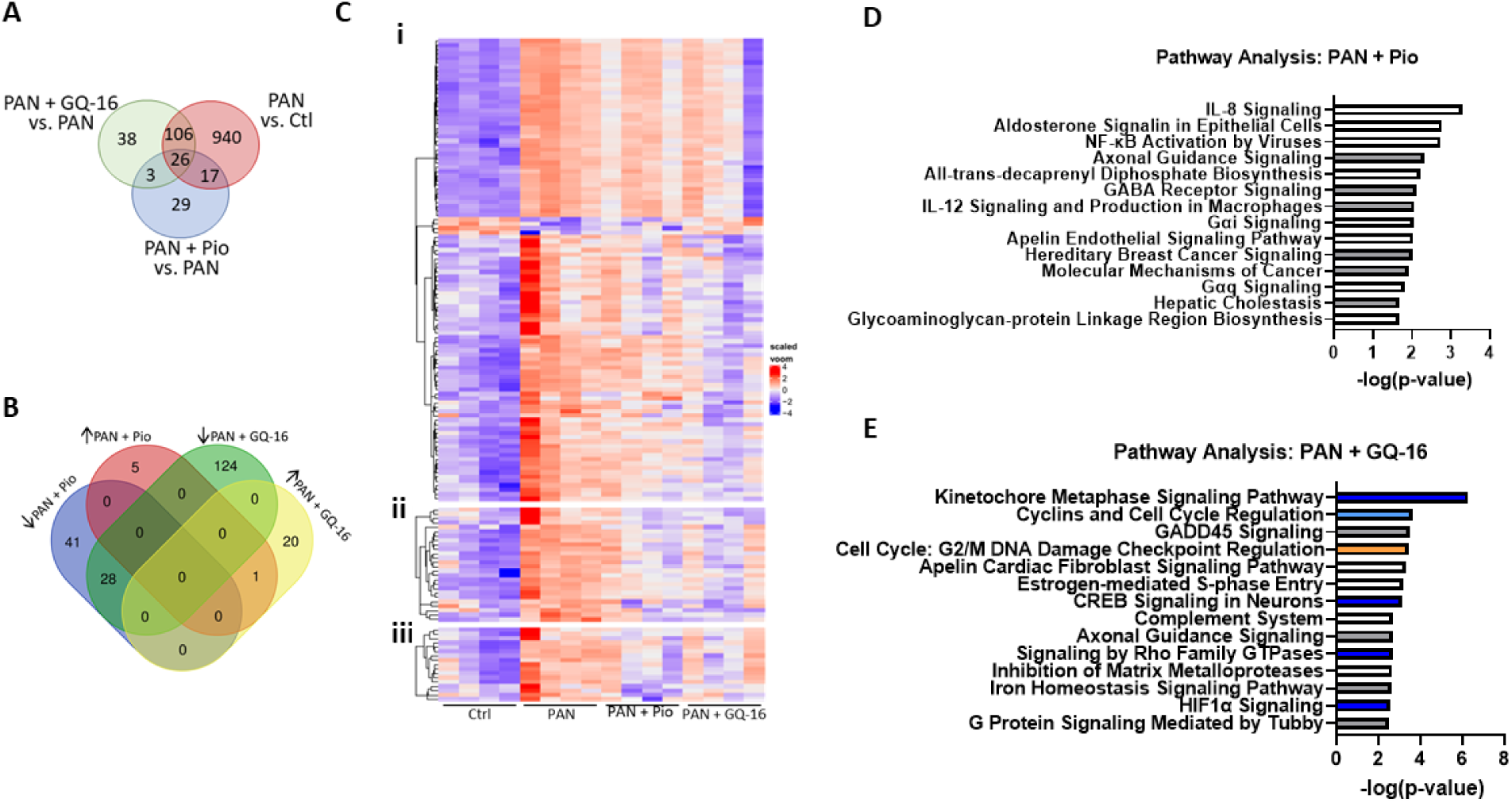
RNASeq Analysis Reveals that Pio and GQ-16 Restore Common and Distinct Glomerular Genes, Pathways and Downstream Processes. **(A)** Differentially expressed genes (DEGs) between PAN vs. Control (pink circle-1089), PAN+GQ-16 vs. PAN (green circle-173) and PAN+Pio vs. PAN (blue circle-75) are depicted. Venn diagrams highlight the number of genes that are commonly (26 DEGs between all 3 groups) or distinctly regulated by the three groups (PAN, PAN+Pio and PAN+GQ-16). **(B)** Venn diagram highlights the DEGs that are up- or down-regulated with Pio and GQ-16 and those that are common and distinct between the two treatment groups, compared to PAN injury. **(C)** Heatmap of DEGs between PAN vs. Control and those restored by (i) only GQ-16, or (ii) both or (iii) only Pio, treatments. Top predicted enriched pathways from Ingenuity Pathway Analysis (IPA) of DEGs between **(D)** PAN+Pio vs. PAN (75 DEGs) and **(E)** PAN vs PAN+GQ-16 vs. PAN (173), are listed and those with positive z-score are denoted in orange, negative z-score in blue, and z score=0 in white.

### Glomerular Disease Associated Hypercoagulopathy is Corrected with GQ-16 Treatment

We have previously shown a significant correlation between hypercoagulopathy and proteinuria during NS, and correction of hypercoagulopathy with glucocorticoid treatment, which is a standard treatment for NS ^39, 40^. We thus measured thrombin generation parameters in our model of NS, their alteration with treatment with Pio and GQ-16, and correlation with proteinuria. A typical thrombin generation curve is shown in Fig. 5A. It is characterized by a short lag phase, the area under the curve (or endogenous thrombin potential), peak thrombin, and velocity index ^42^. Representative curves from each of the study groups are shown in Fig. 5B. ETP is a consistently elevated thrombin generation parameter in NS ^39, 40^, and we found it to be significantly increased in nephrotic rats (PAN, 3646 ± 199 nM*min vs. Control, 2445 ± 402 nM*min; *p* = 0.014) (Fig. 5B and C). This increase in ETP with PAN showed significant reduction with GQ-16 treatment (2544 ± 489 nM*min; *p* = 0.049), while Pio treatment did not have a detectable effect (3509 ± 428 nM*min; *p* = 0.77) (Fig. 5B and C). Notably, ETP strongly correlated with proteinuria (*p* = 0.01) in the nephrotic and treatment rats combined (Fig. 5D). In addition to ETP, other parameters such as peak thrombin generation, lag phase, and velocity index were also derived from these thrombin generation assays (Table 3). The time of the lag phase was significantly reduced with PAN compared to Control, and it showed a tendency to reverse back toward Control with GQ-16 treatment (Table 3).

**Figure 5.**
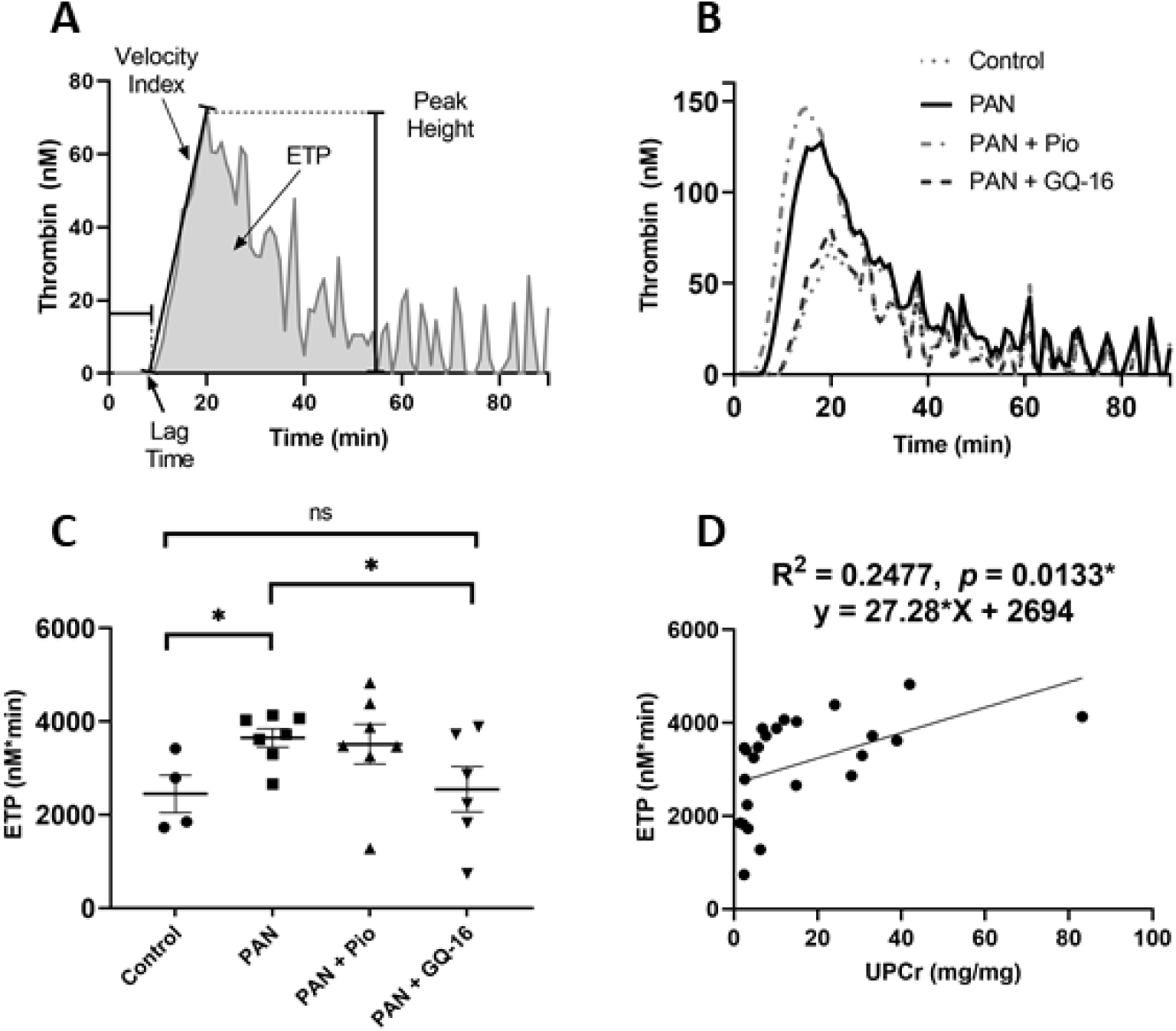
Glomerular Disease Associated Hypercoagulopathy is Corrected with GQ-16 Treatment. Endogenous thrombin potential (ETP) was measured using a thrombin generation assay (TGA) on plasma from rats in each group of PAN-injured, and Pio and GQ-16 treated rats (n=4−7/group). **(A)** Typical thrombin generation curve with ETP, peak height, velocity index, and lag time are labeled. **(B)** Representative graphs of thrombin generation from one rat from each group. **(C)** Mean±SEM of ETP plotted with the individual values shown. **(D)** Linear regression analysis to correlate proteinuria with ETP. * *p* <0.05.

**Table 3.** Coagulation and Thrombin Generation Parameters.

| **Parameters** | **Control** | **PAN** | **PAN + Pio** | **PAN + GQ-16** | **UPCr**<br>**Correlation**<br>**R<sup>2</sup>** | **UPCr**<br>**Correlation**<br>***p* value** |
| --- | --- | --- | --- | --- | --- | --- |
| **ETP**<br>**(nM\*min)** | 2445±402.1 | 3646±198.8*<br>(*p* = 0.0143) | 3509±427.8 | 2544±489.0#<br>(*p* = 0.0493) | 0.2477 | 0.0133± |
| **Peak Thrombin**<br>**(nM)** | 98.97±20.82 | 180.4±27.02<br>(*p* = 0.0698) | 180.8±29.50 | 134.0±35.54 | 0.0282 | 0.4328 |
| **Velocity Index**<br>**(nM/min)** | 16.08±5.073 | 33.07±8.570 | 34.67±8.448 | 19.91±6.643 | 0.0037 | 0.7775 |
| **Lag Phase**<br>**(min)** | 10.38±0.826 | 6.321±0.713**<br>(*p* = 0.006) | 7.714±0.6974 | 9.250±1.532<br>(*p* = 0.0956) | 0.2035 | 0.0269± |

\**p*<0.05; \*\**p*<0.01; compared to Control

### GQ-16 Treatment Reduces Glomerular Disease-Associated Hypercholesterolemia and Alters Hepatic Gene Expression

Dyslipidemia is a major feature of NS and glomerular disease ^8^, manifesting as hypercholesterolemia. In order to understand the role of PPARγ agonists in altering dyslipidemia and the expression of genes involved in lipid metabolism, we measured total cholesterol levels in the serum and the expression of relevant genes in the liver of nephrotic and treated rats. PAN-induced nephropathy resulted in significant increase in total cholesterol levels compared to control (PAN, 161.6 ± 37.6 mg/dL vs. Control, 81.7 ± 2.7; *p* = 0.006) (Fig. 6A). GQ-16 and Pio treatments resulted in significant mean reduction of PAN-induced hypercholesterolemia to 86% (93.1 ± 13.4 mg/dL; *p* = 0.019) and 80% (97.9 ± 12.4 mg/dL; *p* = 0.01), respectively. Moreover, serum cholesterol levels strongly correlated with proteinuria (P<0.0001) in the nephrotic and treatment rats combined (Fig. 6B). *Pcsk9* (encoding for proprotein convertase subtilisin/kexin type 9) expression in the liver tissue was significantly upregulated with PAN-induced nephropathy (∼30 fold) and restored to control levels with both GQ-16 and Pio treatments (Fig. 6C). While *Abca1* (encoding for ATP binding cassette subfamily A member 1) expression was unchanged with PAN, it was increased ∼23 fold with GQ-16 treatment (Fig. 6D).

**Figure 6.**
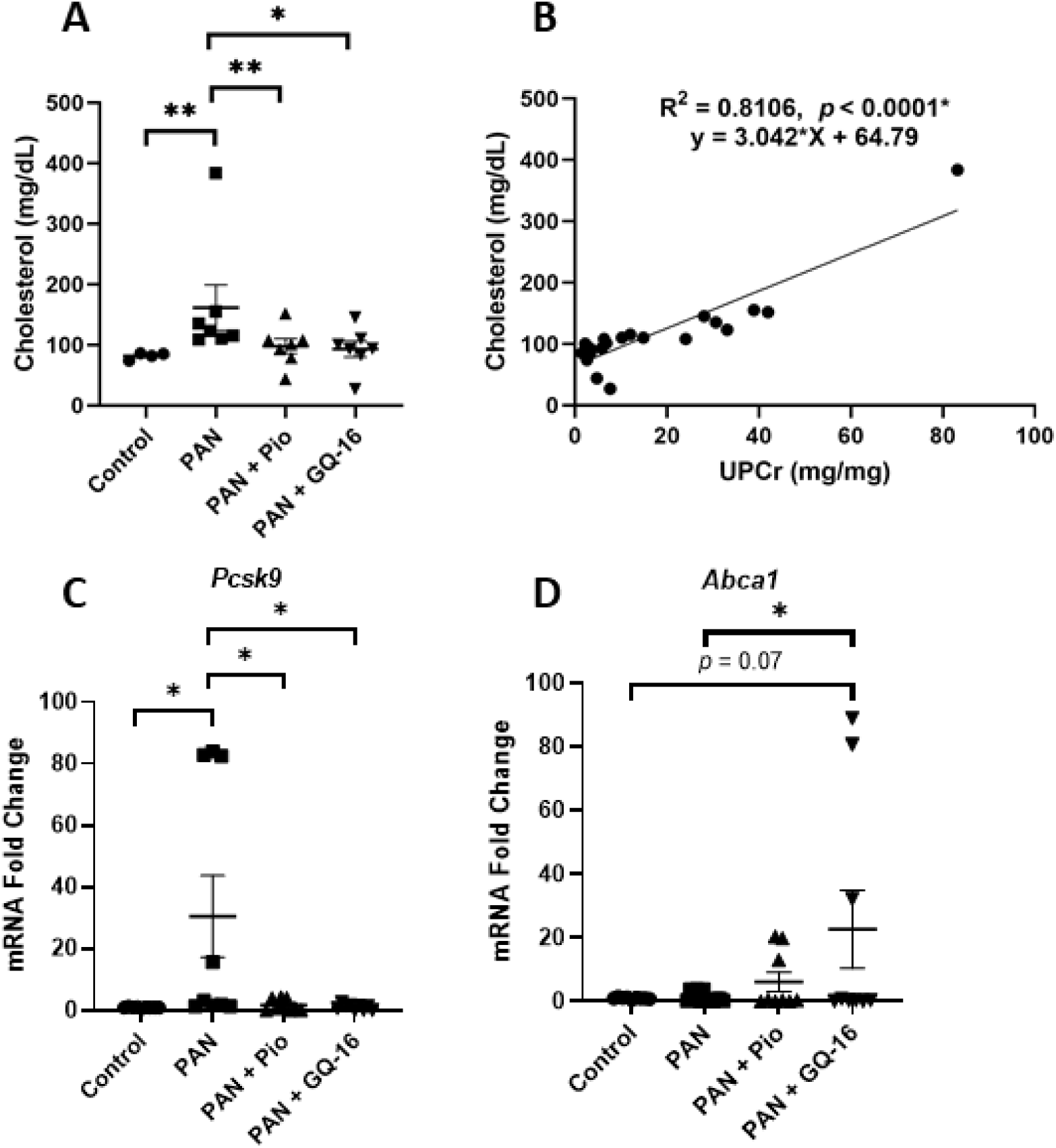
GQ-16 Treatment Reduces Glomerular Disease-Associated Hypercholesterolemia and Alters the Hepatic Gene Expression. Serum cholesterol concentration was measured from Day 11 serum samples and **(A)** plotted as Mean ± SEM. **(B)** Linear regression correlating serum cholesterol with proteinuria. Hepatic gene expression was measured by real time RT-PCR from total RNA extracted from liver tissue from Control, PAN-injected, and PAN-injected rats treated with Pio and GQ-16 (n ≥ 3/group, assay in triplicates). **(C)** *Pcsk9* and **(D)** *Abca1* gene expression was normalized to liver housekeeping gene *Ppia*. Mean ± SEM plotted; * *p* <0.05, ** *p* <0.01.

As liver is also the primary source of albumin (*Alb*) and coagulation proteins, such as prothrombin (*F2*) and antithrombin (*SerpinC1*), and we have observed hypoalbuminemia and hypercoagulopathy with PAN injury and their respective corrections with GQ-16, we measured the hepatic gene expression in the nephrotic and treatment rats (Supplementary Fig. S4). While *Alb* was not induced with PAN, both F2 and SerpinC1 showed a tendency to be induced with PAN and reduced with GQ-16 (*p* = 0.07).

### GQ-16 and Pio Treatments Distinctly Alter Adipogenesis and Gene Expression of Adipogenic Pathways

Adipogenesis and adipogenic pathways are typically induced with PPARγ activation and we thus measured alterations in lipid accumulation and the expression of genes involved in these pathways in differentiated adipocytes *in vitro* and in the epididymal fat of WAT in vivo with Pio and GQ-16 treatments. While both Pio and GQ-16 induced lipid accumulation in differentiated adipocytes in a dose-dependent manner, Pio induced significantly much higher lipid accumulation compared to GQ-16 (Fig. 7Ai). In accordance, expression of *Ap2* (*Fabp4*, encoding for fatty acid binding protein 4), which causes weight gain, was more pronounced and significant with Pio as compared to GQ-16 in both differentiated adipocytes (Fig. 7Aii) as well as in WAT (Fig. 7Bi). On the other hand, expression of insulin sensitizing or secreting adipokines, *Adipoq* (adiponectin) and *Adipisn* (*Cfd*, complement factor D) was induced more robustly with GQ-16 as compared to Pio (Fig. 7B ii and iii). Although a trend was observed towards an increase in *Cd36* (fatty acid transporter) expression, with both Pio and GQ-16, no significant differences were observed (Fig. S5). Notably, as opposed to the studies involving PPARγ in the context of diabetes and obesity, it is not feasible to study the direct effect of weight gain in the current study as PAN-induced treatment itself leads to a modest reduction in weight gain in these rats and limited duration of the study. Nevertheless, we observed that while the weight reduction in nephrotic rats was maintained with GQ-16 treatment, Pio treatment did not show any significant difference from control rats (Supplementary Fig. S6).

**Figure 7.**
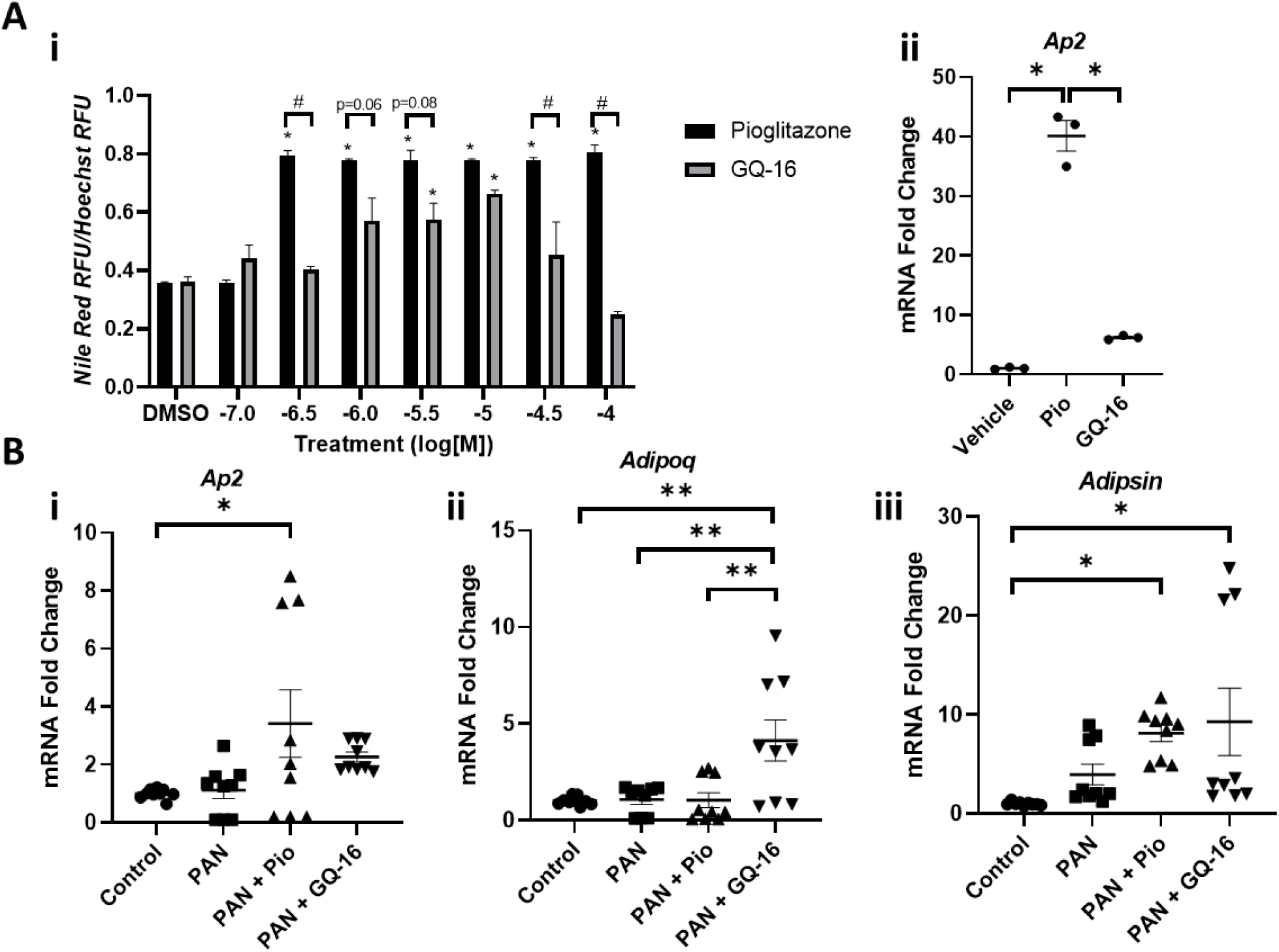
GQ-16 and Pio Treatments Distinctly Alter Adipogenesis and Gene Expression of Adipogenic Pathways. **(A)** 3T3-L1 preadipocytes were cultured and differentiated and assessed for **(i)** lipid accumulation (* p < 0.05 vs DMSO, # p < 0.001) and **(ii)** gene expression of *AP2/FABP4,* in response to Pio and GQ-16 treatments **(B)** White adipose tissue gene expression was measured by real time RT-PCR from total RNA extracted from epididymal fat tissue from Control, PAN-injected, and PAN-injected rats treated with Pio and GQ-16 (n ≥ 3/group, assay in triplicates). Expression of mRNA from **(i)** *Ap2,* **(ii)** *Adipoq,* and **(iii)** *Adipsin* was determined and normalized to the fat house-keeping gene *Ppia*. Mean ± SEM plotted and * *p* <0.05, ** *p* <0.01, *** *p* <0.001, **** *p* <0.0001.

### GQ-16 Treatment Induces Pparg Expression and Reduces Δ Exon-5 Splice Variant Form

While PPARγ co-activator 1α (*Pgc1α*) showed a trend in increase with Pio and GQ-16 treatments (Fig. 8A), both of these treatments induced the expression of *Pparg* (Fig. 8B), which seemed to be more prominent in WAT than in the glomeruli (Fig. 8B and Fig. 3G). Additionally, recently a naturally spliced variant form of Pparg has been identified in human adipocyte progenitor cells and in WAT of obese and diabetic patients ^43^. We were able to detect the expression of a Δ exon 5 splice variant form of *Pparg*, which was found to be decreased with injury as well as with both Pio and GQ-16 treatments in the WAT when normalized to the total *Pparg* expression (Fig. 8C). Notably, GQ-16 further reduced the levels of Δexon 5 *Pparg* variant.

**Figure 8.**
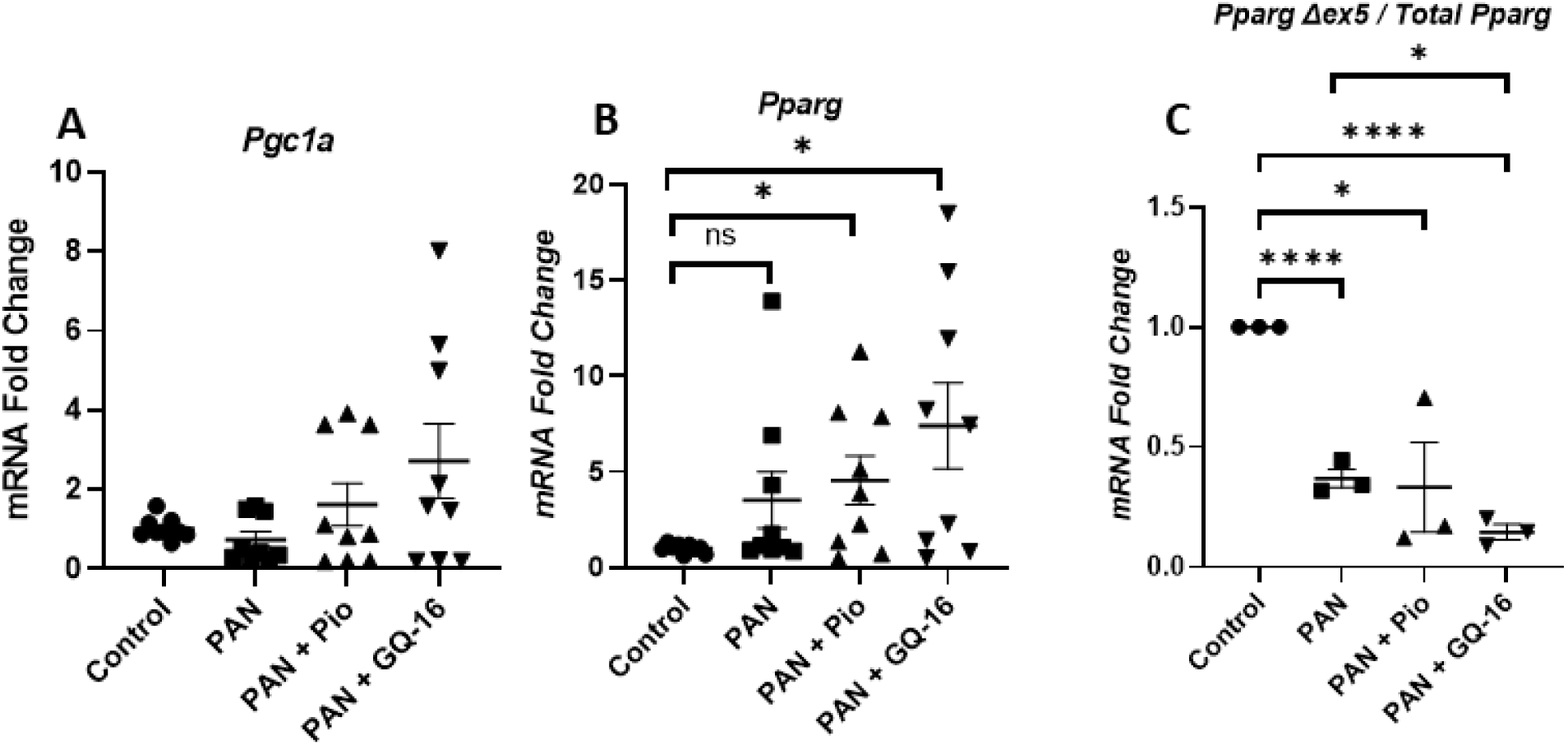
GQ-16 Treatment Induces *Pparg* Expression and Reduces Δ Exon-5 Splice Variant Form. White adipose tissue gene expression was measured by real time RT-PCR from total RNA extracted from epididymal fat tissue from Control, PAN-injected, and PAN-injected rats treated with Pio and GQ-16 (n ≥ 3/group, assay in triplicates). Expression of mRNA from **(A)** *Pgc1a,* and **(B)** *Pparg* was determined and normalized to the fat house-keeping gene *Ppia*. **(C)** Expression of *Pparg* Δ exon 5 splice variant form was measured and normalized to *Pparg.* Mean ± SEM. * *p* <0.05, ** *p* <0.01, *** *p* <0.001, **** *p* <0.0001.

## Discussion

Glomerular disease is the leading cause of ESKD in the United States, and NS, characterized by high-grade proteinuria, is one of the most common forms of glomerular disease ^1–3^. Furthermore, NS is typically associated with hypoalbuminemia, hypercholesterolemia, systemic immune dysregulation, hypercoagulopathy, and edema ^4–8^. Standard treatments include glucocorticoids (for idiopathic and primarily pediatric NS), but they have many side effects, and 10-50% of adult and pediatric NS patients can be resistant to steroid treatment ^44–46^. NS patients may exhibit hypertension, which can be managed with ACE inhibitors and ARBs, which reduce proteinuria. Moreover, steroid-resistant NS is associated with an increased risk of developing CKD, which account for 15% of all children with CKD requiring renal replacement therapy ^47^. Thus, there is an urgent and critical need to develop new alternative therapies for NS and glomerular disease with increased efficacy and reduced side effects. In order to do so, we and others have previously reported that PPARγ agonists not only provide beneficial protective effects in type II diabetes and DN, but also in various models of non-diabetic glomerular disease ^13–21^. However, targeting PPARγ via widely marketed anti-diabetic drugs (traditional TZDs) has been re-evaluated due to significant side effects such as weight gain, heart failure, bone fracture, and bladder cancer ^30–35^. In the current study, we hypothesized that the downstream effects of PPARγ can be mechanistically dissociated and that selected modulation of PPARγ by a novel partial agonist, GQ-16, will result in a better and targeted therapeutic advantage in glomerular disease. Our results revealed that GQ-16 is highly efficacious in reducing proteinuria in a PAN-induced animal model of glomerular disease or NS (Fig. 9). Our results further demonstrated that while both GQ-16 and Pio restored glomerular *Nphs1* and hepatic *Pcsk9* expression and reduced hypercholesterolemia, the beneficial effects of GQ-16 were also associated with restoration of glomerular *Nrf2*, and reduction in disease-associated hypoalbuminemia and hypercoagulopathy. Furthermore, RNA-seq analysis identified both common and distinct glomerular genes and pathways altered by Pio and GQ-16 treatments. Moreover, both in adipocytes and in WAT, Pio but not GQ-16 treatment resulted in significant induction of *Ap2* (fatty acid binding protein). Furthermore, Pio induced significantly more lipid accumulation in differentiated adipocytes compared to GQ-16. Additionally, both Pio and GQ-16 induced insulin sensitizing adipokines in WAT with varying degrees. Taken together, these findings suggest that PPARγ can be selectively modulated by its partial agonist GQ-16, in order to provide the beneficial proteinuria-reducing effects and reduce NS associated co-morbidities, while reducing the side-effects conferred by traditional PPARγ full agonists.

**Figure 9.**
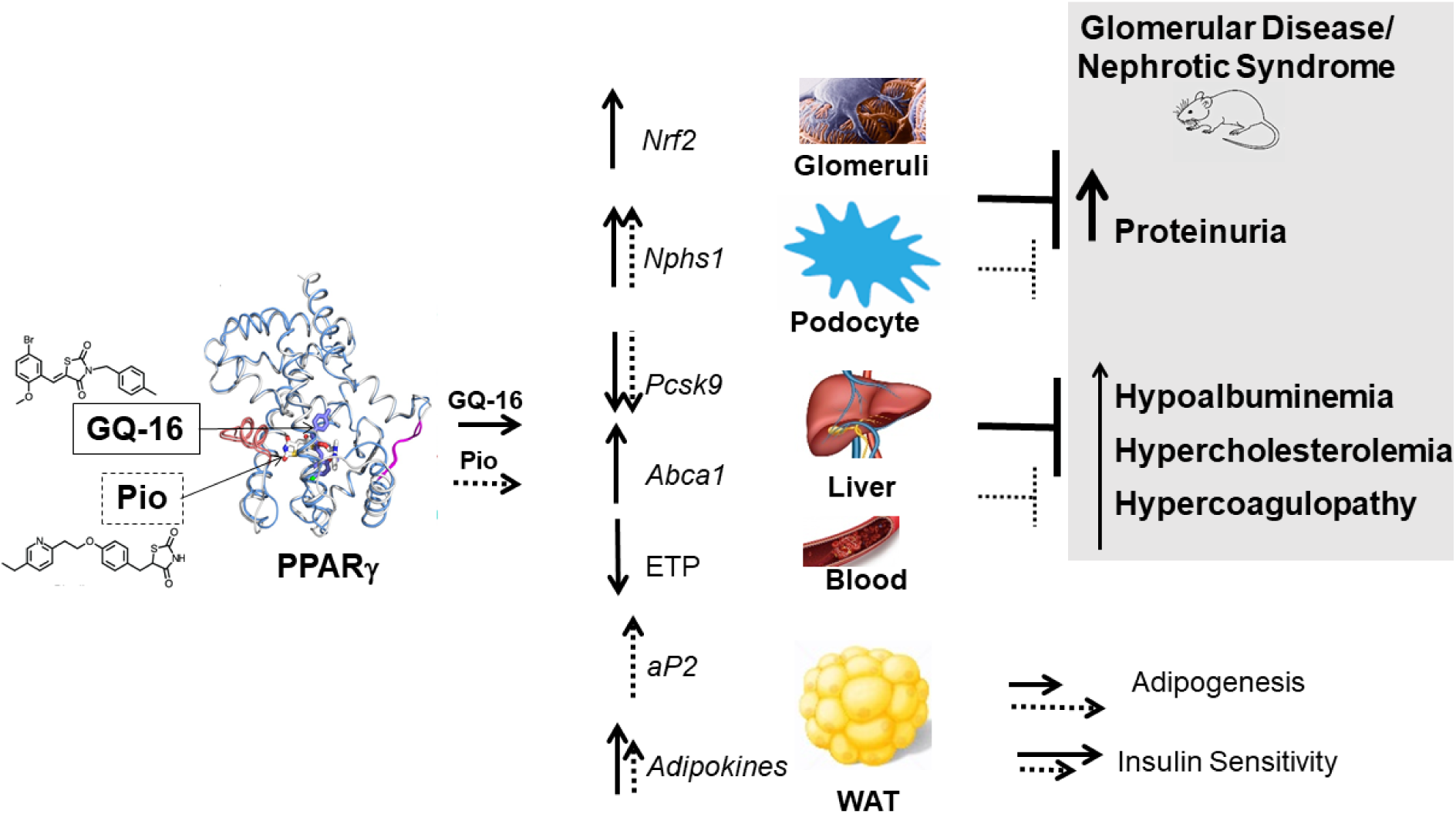
Schematic Depicting the Beneficial Effects of GQ-16 in Improving Glomerular Disease. The traditional full agonist of PPARγ, pioglitazone (Pio) and the partial agonist and selective modulator, GQ-16, both bind and activate PPARγ, although in distinct ways. While GQ-16 activates PPARγ differently than Pio, it is equally or more efficacious in reducing proteinuria and overall nephrotic syndrome-associated co-morbidities such as hypoalbuminemia, hypercholesterolemia and hypercoagulopathy. These effects are associated with increased glomerular *Nrf2* expression, increased expression of *Nphs1* in podocytes, decreased hepatic expression of *Pcsk9* and increased hepatic *Abca1* expression, and reduced endogenous thrombin potential (ETP) in the plasma of nephrotic rats treated with GQ-16. Moreover, GQ-16 induces adipocyte lipid accumulation and white adipose tissue (WAT) *aP2* to a lesser extent than Pio and it increases the expression of adipokines to a larger extent than Pio, which likely renders reduced adipogenesis and increased insulin sensitivity as compared to Pio. Solid arrows and lines represent GQ-16 effects and dashed arrows and lines represent the effects of Pio.

PPARγ is attributed to the therapeutic basis of TZDs to treat diabetes, since it improves insulin sensitivity and decreases insulin demands ^32^. While PPARγ is the master regulator of glucose and lipid metabolism and regulator of adipogenesis, in the last couple of decades, various preclinical studies and meta-analyses have highlighted its direct beneficial role in kidney cells in addition to its favorable systemic effects in the context of diabetes ^48, 49^. Notably, in the last decade, we and others have documented the beneficial roles of TZDs in directly protecting podocytes from injury ^9–12^, in reducing proteinuria and glomerular injury in various animal models of glomerular disease such as minimal change disease, focal segmental glomerulosclerosis (FSGS), and crescentic glomerulonephritis^13–19, 21^, and in improving clinical outcomes in NS patients refractory to steroid treatment ^13, 22^. More recently, breakthrough discoveries in the field of PPARγ biology suggest that the beneficial insulin-sensitizing activities of PPARγ can be dissociated from its harmful adipogenic activities ^28, 29, 36, 50, 51^. This has led to the development of mechanistically distinct novel compounds, including GQ-16, that have weak traditional PPARγ agonistic activity (adipogenic) and associated weight gain effects, but very good anti-diabetic activity ^37, 38^.

In the present study, we have specifically demonstrated that GQ-16 reduces proteinuria in a rat model of NS with great efficacy. The 84% reduction in proteinuria by GQ-16 compared to PAN-nephrosis, was found to be comparable to controls as well as to the previously described reduction with high dose glucocorticoids (79%) ^13^. GQ-16 treatment was also effective in correcting another characteristic feature of NS, hypoalbuminemia in PAN NS. Furthermore, podocyte injury and loss are characteristic features of proteinuria in NS and the integrity of podocyte signaling pathways and slit diaphragm is paramount to their proper structure and function ^52^. Nephrin is a critical podocyte marker and an essential component of the slit diaphragm, which is either mutated in the congenital form of NS (Finnish Type) or decreased in expression in various forms of glomerular disease ^13, 52–54^. In accordance with its essential role in glomerular function, we found that *Nphs1* expression inversely correlated with proteinuria and it was significantly reduced in the glomeruli of the nephrotic rats and increased with treatments with PPARγ agonists. NRF2 is the master regulator of oxidation pathways and is well known to have cytoprotective and anti-inflammatory effects ^55^. It also plays a critical role in regulating PPARγ expression ^56^ and it’s complete deletion has recently been demonstrated to result in decreased podocyte PPARγ and an aggravated course of glomerular disease as reported in a rapid proliferative glomerulonephritis model ^14^. Moreover, enhanced podocyte injury seen in the animal models of glomerular disease with podocyte specific *Pparg* deletion is also associated with reduced expression and activity of NRF2. Accordingly, we found a significant decrease in the expression of glomerular *Nrf2* in nephrotic rats, which was only mildly restored with Pio, but completely completely by GQ-16 to control levels. Gain-of-function mutations in TRPC6 and its increased podocyte expression and activity is linked with glomerular injury and proteinuria, especially in chronic models ^57, 58^. While, it has been demonstrated it may act as a mediator of sildenafil-mediated protection of podocyte injury by inhibition via PPARγ in an adriamycin-induced focal segmental glomerulosclerosis (FSGS) ^20, 21^, we did not observe a significant increase during PAN nephrosis and decrease in its expression with PPARγ agonists. Our studies are in alignment with previous observations where enhanced TRPC6 activation by mechanical stimuli is observed in FSGS, but not in acute phase of PAN nephrosis and in contrast to the chronic model, *Trpc6* del/del rats are not protected during the acute phase of PAN nephrosis ^59^. Furthermore, it is clear from the glomerular transcriptome data and analysis that the beneficial effects of Pio and GQ-16 treatments are associated with elicitation of both common and distinct glomerular DEGs, pathways and biological processes. These results together suggest that GQ-16 is equally as or perhaps more efficacious than Pio in reducing glomerular injury and proteinuria, while eliciting common and distinct downstream glomerular events and selectively modulating PPARγ.

PPARγ has been shown to be phosphorylated in obesity models at the Ser273 site and dephosphorylated with the insulin sensitizing effects of PPARγ agonists ^28, 29, 36^. Moreover, in an *in vitro* assay, GQ-16 has been shown to dephosphorylate PPARγ like traditional TZDs ^37, 38^. Interestingly, our results demonstrate that relative phosphorylated PPARγ levels (at Ser273) remain unaltered with PAN injury and subsequent Pio and GQ-16 treatments. This result suggests that while PPARγ-Ser273 is known to play a major role in determining the insulin sensitizing effects of PPARγ in adipose tissue, the same mechanism may not play a role in determining its proteinuria-reducing effects in glomeruli.

Our previous observations have found increased ETP to be very well-correlated with increased proteinuria during NS and its decrease with standard treatment in both human studies and in animal models ^39, 40^. In the current study, we find a good correlation of ETP with proteinuria, significant increase with injury, and a decrease with GQ-16 treatment. Earlier study from our group demonstrated that Pio treatment decreased ETP in concert with proteinuria during PAN NS. However, Pio significantly increased ETP when given to healthy control rats, suggesting its inherent role in increasing ETP ^40^. In the current study, we failed to observe a significant reduction in ETP with Pio in NS, which could be because of Pio’s inherent ability to increase ETP. Additionally, Pio used in the present study was from a different manufacturer, and the time between the last Pio dose and PPP collection was decreased by approximately 8 hours. This likely contributed to increased levels of Pio in the blood at the time of ETP measurement, as the systemic pharmacokinetic profile of Pio suggests that it peaks at 1.5-2 hours and its half-life is approximately 8-9 hours ^60, 61^. Overall, our studies suggest that partial agonism of PPARγ with GQ-16 provides a better correction of hypercoagulopathy, than full agonism by Pio.

Dysregulated lipid metabolism is one of the other major features of NS and glomerular disease ^8^. Hepatic levels of PCSK9 are known to be upregulated in glomerular disease which contributes to dyslipidemia by degrading the low density lipoprotein (LDL) receptor and decreased LDL uptake by the liver ^8^. Inhibition of PCSK9 using anti-PCSK9 monoclonal antibodies and siRNAs has gained high clinical importance due to its lipid-lowering effects in the conditions of hypercholesterolemia and is gaining traction in the context of NS with the emergence of case reports and new clinical trials ^8, 62–65^. Our results corroborate the importance of PCSK9, as we found a significant induction in its gene expression in the liver of nephrotic rats. Moreover, both Pio and GQ-16 robustly and significantly reduced the PAN-induced hepatic *Pcsk9* expression to control levels. ABCA1 plays an important role in cholesterol efflux and experimental evidence in animal models of FSGS suggests that an increase in ABCA1 activity mainly in the liver and other peripheral organs, including in podocytes, is associated with improved proteinuria ^8, 66–69^. Interestingly, while the increase was variable, we observed a significant increase in its expression with GQ-16 treatment, but only a trend with Pio in our animal model of NS.

GQ-16 has been shown to exhibit modest adipogenic activity when compared to TZDs as shown by reduced transactivation activity and *Ap2* induction ^37, 38^. Findings in obese Swiss male mice indicated that in addition to exhibiting insulin-sensitizing properties, 14-day treatment with GQ-16 induced decreased edema, weight gain, and visceral WAT mass in response to high fat diet, despite increasing energy consumption ^37, 38^. These effects were accompanied by the induction of thermogenesis-related genes in epididymal fat depots, suggesting that browning of visceral WAT may have contributed to weight loss. These results strongly support that PPARγ activation by partial agonists, devoid of full agonism-related unfavorable effects, may be a strategy to induce browning of WAT and hence to treat obesity and diabetes. Specifically, our current study supports this theory in the context of non-diabetic disease. Our studies show that Pio induces increased lipid accumulation and *Ap2* induction as compared to GQ-16 in differentiated adipocytes. Similar induction of *Ap2* was also observed in the WAT epididymal fat of Pio-treated rats, but not in GQ-16 treated rats. Furthermore, both full and partial PPARγ agonists are known to increase the expression of insulin sensitizing adipokines, adiponectin and adipsin, as well as the fatty acid transporter CD36 ^36^. While we did not observe a significant change in the expression levels of CD36, both Pio and to a larger extent GQ-16, induced *Adipsin* and *Adipoq* expression. Overall, our results suggest that GQ-16 has a lower adipogenic profile than Pio, which would potentially offer a therapeutic advantage during long tern treatment.

PPARγ exists in mainly two major isoforms, γ1 and γ2, which are a result of different promoter usage as well as alternative splicing (Fig. 1) ^20, 26, 27^. While γ1 is ubiquitously expressed, including in podocytes, γ2 is mostly restricted to adipose tissue and liver (data not shown). Moreover, the recently identified Δ exon-5 splice variant form of *Pparg* has been shown to function as a dominant negative form by reducing the adipogenic potential of precursor cells and its increased expression in the adipose tissue has shown positive correlation with body mass index in obese and diabetic patients ^43^. We have observed the presence of a Δ exon5 spliced form of *Pparg* in the WAT, which was reduced with PAN injury and further decreased with GQ-16 treatment. Although speculative and open to interpretation as decrease in Δexon5 form is minor and relatively enhanced by an increase of total *Pparg* form, it is possible that a decrease in Δexon5 *Pparg* with PAN injury could be an adaptive mechanism, and its further reduction by GQ-16 in our study serves as a positive feedback loop to enhance PPARγ activity. Moreover, we were unable to detect a robust and consistent expression of the Δexon5 form in the podocytes.

The current study has some limitations and strengths. While we have demonstrated the efficacious and beneficial role of GQ-16 in PAN-nephrosis, future studies in other non-diabetic models of glomerular disease, such as chronic PAN model, adriamycin-induced FSGS, or even immune-mediated forms of glomerular disease could provide a broader treatment potential for GQ-16. Moreover, future research which is out of the scope of the current study, is warranted to enhance the clinical appeal of the beneficial effects of GQ-16, which would demonstrate its effects after disease onset. Nevertheless, our study provides a proof of concept for protection of experimental NS with an existing novel compound and PPARγ modulator and opens up new avenues of designing or adapting new modulators of PPARγ for the treatment of glomerular disease. Furthermore, while we have measured dose-dependent effects and drawn comparisons of GQ-16 and Pio in PPARγ activation, lipid accumulation, and *Ap2* induction, and demonstrated decreased PPARγ activation with a lower adipogenic profile of GQ-16 compared to Pio, our *in vivo* studies are limited by a single dose study. However, we have carefully chosen the *in vivo* doses for the two compounds, based on the *in vitro* results and previous *in vivo* studies in which we observed protective effects of Pio in NS^13^ and reduced adipogenic profile of GQ-16 ^38^. While it is not feasible to study weight gain in a short-term nephrosis model, a real clinical scenario would benefit from long term treatment with a compound which offers reduced weight gain potential. The strength of our study is that GQ-16 has equal or higher efficacious proteinuria-reducing effects than Pio, reduction in NS-associated co-morbidities while providing reduced adipogenic effects, as measured by reduced *Ap2* induction in vitro and in vivo and lipid accumulation in vitro as compared to Pio.

In summary, our studies suggest that selective modulation of PPARγ by a partial agonist is efficacious in reducing proteinuria, and is perhaps more beneficial than a full PPARγ agonist in reducing hypoalbuminemia and hypercoagulopathy, while providing reduced adipogenic potential and drug-induced side effects in a PAN-induced animal model of NS. Our findings not only emphasize the benefits of GQ-16 as a novel therapeutic modality for NS and deepen our molecular understanding of the role of PPARγ in glomerular disease, but open up new possibilities and potential future clinical implications for selectively modulating PPARγ by partial agonists for the treatment of glomerular disease. In this regard, designing novel modulators of PPARγ which would result in its optimal conformation that would yield desirable molecular and clinical effects, would be highly significant.

## Acknowledgments

We thank Jeffrey Miner, Washington University, St Louis, Missouri for his valuable comments and suggestions and for his mentorship on the American Heart Association Career Development Award granted to SA. We also thank Alexander S Banks, Beth Israel Deaconess Medical Center, Harvard Medical School, Boston, MA for critical comments and review of the manuscript. We thank the Genomics Core Resources at The Ohio State University for RNA Sequencing Services.

## Data Sharing

The RNAseq data is available on GEO with accession GSE179945 (https://www.ncbi.nlm.nih.gov/geo/query/acc.cgi?acc=GSE179945) “token mdoxmokgrnodpex for the Reviewers”

## Author Contributions

CB performed experiments, analyzed and interpreted the data, prepared figures and tables, and drafted the manuscript. GR, APW and AAA performed experiments and edited the manuscript. BAK and APW interpreted the coagulopathy data and edited the manuscript. RC performed the histological analysis and edited the manuscript. MRG synthesized and provided the compound GQ-16 for these studies and edited the manuscript. AW performed bioinformatics analysis and edited the manuscript. FARN, BB and AF interpreted the data and edited the manuscript. SA conceptualized and designed the study, analyzed and interpreted the data, prepared figures and tables, and drafted and edited the manuscript. All the authors approve of the final version of the manuscript. Part of these findings have been selected for Platform Presentation at the 13^th^ International Podocyte Conference to be held at the University of Manchester, July 27-31, 2021.

## Supplementary Material

### Methods

#### Animal Studies: Glomerular Disease Model and Treatment with GQ-16 and Pio

This study was approved by the Institution Animal Care and Use Committee at Nationwide Children’s Hospital, and their guidelines were followed when performing the experiments. GQ-16 was synthesized as previously described and quality tested for 99% purity ^1^. Male Wistar rats were intravenously (IV) injected with puromycin aminonucleoside (PAN) (Sigma-Aldrich, St. Louis, MO) (50 mg/kg) on Day 0, which induced proteinuria, while control rats were given IV injections of saline. The rats were then treated by oral gavage with Pioglitazone (Alfa Aesar, Tewksbury, MA) (10 mg/kg), GQ-16 (40 mg/kg), or a sham vehicle daily. GQ-16 and Pio dosages were determined based on in vitro PPARγ activation, *aP2* induction and lipid accumulation data in the current study and our previous studies on the expression of thermogenesis-related genes and adipogenic effects of GQ-16 ^2^ and proteinuria reducing beneficial effects by Pio ^3^. Spot urine and serum were collected, and body weights were recorded throughout the study. The rats were anesthetized with 3% isoflurane and sacrificed on Day 11, at which time blood was collected through the inferior vena cava with a 23-G needle into the 0.32% sodium citrate and 1.45μM corn trypsin inhibitor, then processed to Platelet Poor Plasma (PPP) as described previously ^4, 5^. Kidneys were harvested, and the glomeruli were isolated from 1 and ½ kidneys using the sequential sieving method ^3^. Half of the kidney was fixed in 10% buffered formalin for histologic evaluation. Cross sections of kidney containing cortex, medulla and papilla were processed routinely, sectioned at 4 µm thickness and stained with periodic acid-Schiff method. Slides were reviewed by a pathologist blinded to the treatment method. Liver and white adipose tissue (WAT) epididymal fat were collected and flash frozen in liquid nitrogen.

#### Urinalysis

Urine was collected from rats daily throughout the study and resolved using sodium dodecyl sulfate-polyacrylamide gel electrophoresis (SDS-PAGE) on an 8% gel and stained with Coomassie Brilliant Blue G-250 (Alfa Aesar, Tewksbury, MA) to visualize the albumin bands. Urine protein:creatinine ratio (UPCr) analyses were performed on urine samples from Day 11 at the time of peak proteinuria by Antech Diagnostics GLP (Morrisville, NC) to quantify the proteinuria values ^3, 4^.

#### Serum Chemistry

Serum albumin and cholesterol were measured using the ACE® albumin and cholesterol reagents (Alfa Wasserman Diagnostic Technologies, LLC, West Caldwell, NJ) on the Vet Axcel (Alfa Wasserman Diagnostic Technologies, LLC, West Caldwell, NJ) at the Clinical Pathology Services, The Ohio State University’s College of Veterinary Medicine, according to the manufacturer’s instructions.

#### Coagulopathy Measurement

Thrombin generation assays (TGA) were performed using the Technothrombin TGA Kit (Technoclone, Vienna, Austria) and Reagent C (RC) Low on PPP samples collected from the rats to determine various parameters such as the endogenous thrombin potential (ETP) and peak thrombin concentration. These assays were performed at least in duplicate on various rat groups (n=4-7/group) as previously described ^4^. Briefly, PPP at 1:1 ratio with buffer was added to black well plates, and RC Low added. Then TGA substrate was added just before reading on a Spectramax M2 Fluorescent Plate Reader (Molecular Devices, San Jose, CA).

#### RNA Isolation, Quantitative Real Time Reverse Transcription-Polymerase Chain Reaction

Total RNA was extracted from isolated WAT epididymal fat and liver tissue samples using the RNeasy kit (Qiagen, Germantown, MD), following the manufacturer’s instructions. Tissue samples in lysis buffer were placed in lock tubes with stainless-steel disruption beads and lysed at 30.0 Hz for 4 minutes using the Qiagen TissueLyser (Germantown, MD), followed by RNA isolation from the resulting lysate. Total RNA was isolated from glomeruli tissue samples using the mirVana Isolation Kit (Invitrogen, Carlsbad, CA), according to the manufacturer’s instructions. Yield and purity were calculated for all isolated RNA samples by measuring the absorbance at 260, 280, and 230 nm and the ratios (both) with a spectrophotometer. 500 ng - 1µg of RNA was subjected to DNase (Invitrogen, Carlsbad, CA) digestion at room temperature for 15 minutes, which was then inactivated with 25mM EDTA at 65°C for 10 min. RNA was reverse transcribed using the iScript cDNA Synthesis Kit (Bio-Rad, Hercules, CA) according to the manufacturer’s instructions. cDNA was used for quantitative reverse transcription-polymerase chain reaction (qRT-PCR) using gene specific and house-keeping primers (Table 1). SYBR green (Bio-Rad, Hercules, CA) qRT-PCR was performed on the Applied Biosystems 7500 Real-Time PCR System. The PCR conditions were 95°C for 10 minutes, 40 X (95°C for 15 seconds, 60°C for 1 minute), followed by a melt curve to ensure specific products. The annealing temperature was at 60°C for all genes except *Pgc1a, Pparg,* and *Cd36* for which 52°C annealing temperature was used. The melt curve conditions were 95°C for 15 seconds, 60°C for 1 minute, 30 seconds incremental increase to 95°C and 60°C for 15 seconds, as described previously ^3^. The ΔΔCt method ^6^ was used to analyze the results, including normalization to housekeeping genes (*Rpl6* ^7^ for glomeruli, *Ppia* ^8, 9^ for fat and liver). Due to melt curve variation for *Trpc6,* amplified samples were also resolved on a 2% agarose gel, and densitometry was performed using ImageJ (National Institutes of Health, Bethesda, MD) software. Each gene was tested in triplicates for each tissue on at least three different rats per group.

#### RNA Sequencing, Differential Gene Expression, Pathway Analysis and Ontology Enrichment

For RNA sequencing, mRNA libraries were generated using ∼200ng total glomerular RNA (quantified using Qubit Fluorometer) with RIN of >7, using NEBNext Ultra II Directional (stranded) RNA Library Prep Kit for Illumina (NEB #E7760L), NEBNext Poly (A) mRNA Magnetic Isolation Module (NEB #E7490) and NEBNext Multiplex Oligos for Illumina Unique Dual Index Primer Pairs (NEB #6442S/L). Libraries were sequenced with Illumina NovaSeq 6000 flow cell using paired-end 150-bp format to at least 17 million passed-filter clusters/sample (equivalent to 34 million reads). Glomerular RNA was subjected to RNA sequencing using NovaSeq6000 SP 300 cycles (=2×150bp) and using internal pipeline ^10^, reads were aligned to Rat genome Rnor6.0 with HISAT2 ^11^ and counts generated for Rnor 6.0 v101 with featureCounts from the subread package ^12^. Post alignment quality check (QC) was assessed with fastqc, RSeQC, and picard ^13^ (https://broadinstitute.github.io/picard). Counts were normalized with voom and differential expression tested with limma ^14^. Differential expressed genes (DEGs) were chosen with p<0.05 and abs(logFC)>1. Heatmaps were generated with ComplexHeatmap in R. Enriched canonical pathways were identified using Ingenuity Pathway Analysis (IPA) and GO term enrichment performed with MOET - Multi Ontology Enrichment Tool (MOET, Ontology Enrichment (mcw.edu)) to identify enriched biological processes, cellular components and molecular functions.

#### Protein Isolation, SDS-PAGE and Western Blotting

To isolate protein from the glomeruli, liver, and WAT epididymal fat tissues, the samples were lysed in a lock tube with RIPA buffer (1M Tris HCl, 0.5M EDTA, 5M NaCl, 10% SDS) containing protease inhibitor cocktail (Thermo Scientific, Waltham, MA) and phosphatase inhibitor cocktail (Alfa Aesar, Tewksbury, MA) and a stainless-steel disruption bead using a Qiagen TissueLyser (Germantown, MD). Following lysis/homogenization at 30.0 Hz for 4 minutes (liver and fat) or 1 minute (glomeruli), samples were centrifuged for 10 minutes at 4°C at 12,000rpm. The supernatant was collected, and the proteins were resolved by SDS-PAGE and transferred to an Immobilon-P polyvinylidene difluoride (PVDF) Transfer Membrane (Millipore Sigma, St. Louis, MO). The membrane was blocked with 5% milk in phosphate buffer saline (PBS) with 0.1% Tween 20 (PBST) for 1 hour, followed by incubation with the primary antibody overnight [anti-Phospho-PPARγ-Ser 273 (Bioss, Woburn, MA), anti-PPARγ (Proteintech, Carlsbad, CA), and anti-GAPDH (Cell Signaling, Danvers, MA)]. The membrane was washed three times in PBST and incubated with the secondary antibody [anti-rabbit IgG (Jackson ImmunoResearch Laboratories, Inc, West Grove, PA)] in 5% milk in PBST for an hour. Protein bands were detected by chemiluminescence using the Chemidoc MP Imaging System (Bio-Rad, Hercules, CA). Densitometry was performed on the bands using ImageJ (National Institutes of Health, Bethesda, MD) software, and band density was subtracted from the background and normalized to GAPDH.

#### PPARγ Activation Luciferase Assay

HeLa cells were transiently co-transfected with plasmids containing human *PPARG* Ligand-Binding Domain fused to the *GAL4* DNA-Binding Domain and a plasmid containing the luciferase reporter gene under the regulation of five Gal4 DNA-binding elements (UASG × 5 TK-luciferase). These plasmids were kindly provided by Dr. Paul Webb from Methodist Research Institute, TX, United States. Transfections were conducted using the Lipofectamine 2000 Transfection Reagent (Invitrogen, Carlsbad, CA), according to the manufacturer’s instructions. Briefly, cells were plated at the density of 25,000 cells per well in 48-well plates and maintained at 37°C and 5% CO_2_ for 24 h. Cells were then transfected with *hPPARG*-LBD (60 ng per well), UASG5x-Luc (240 ng per well), and pCMV-β-galactosidase (60 ng per well, used as an internal control). After 6 h, the culture medium was replaced by fresh medium containing DMSO (vehicle control) or increasing concentrations of Pio or GQ-16 (0.1 nM to 100 μM). Luciferase activity was measured using the Luciferase Assay System Kit (Promega, Madison, WI) in a luminometer (GloMax^®^ 20/20 Luminometer - Promega, Madison, WI), following manufacturer’s instructions. Results were reported as mean luciferase activity induced by the different ligands relative to vehicle control. Each experiment was performed in triplicate and repeated at least three times.

#### 3T3-L1 Preadipocyte Culture and Differentiation, Lipid Accumulation Assessment and aP2 Induction

3T3-L1 preadipocytes were obtained from the cell bank of the Federal University of Rio de Janeiro (Rio de Janeiro, Brazil) and expanded at 37°C and 5% CO_2_ in DMEM supplemented with 10% calf bovine serum, 100 IU/mL Penicillin, and 100 μg/mL Streptomycin. Cells were plated in 6-well plates for gene expression analysis (48 × 10^3^ cells/well) or in 24-well plates for intracellular lipid accumulation assessment (12 × 10^3^ cells/well). Two days after confluence, culture medium was switched to DMEM supplemented with 10% fetal, 100 IU/mL Penicillin, and 100 μg/mL Streptomycin, and adipose induction cocktail containing 0.5 mM isobuthylmethylxnathine (Sigma-Aldrich, St Louis, MO), 250 nM dexamethasone (Sigma-Aldrich, St Louis, MO) and 1 μg/mL insulin (Sigma-Aldrich, St Louis, MO). After 72 hours, cells were maintained with DMEM containing 1 μg/mL insulin. For intracellular lipid accumulation assessment, cells were plated in 24-weel plates (12 × 103 cells/well), and induced as described above, but with removal of isobuthylmethylxnathine from induction medium. Vehicle control (0.1% DMSO) or ligands (100 μM pioglitazone or 100 μM GQ-16) were added throughout all adipose differentiation period. Culture medium was changed every three days, and cells were harvested 5 days after adipose induction for RNA isolation or after 10 days for lipid accumulation assessment. All experiments were conducted in triplicate.

#### Lipid Accumulation Assessment

3T3-L1 preadipocytes induced to differentiate into adipocytes were fixed in 3.7% formaldehyde. Intracellular neutral lipids were stained with Nile Red (1 μg/mL), and nucleic acid was stained with Hoechst 33342 (5 μg/mL). Relative fluorescence units (RFUs) were measured in EnSpire Multimode Plate Reader (PerkinElmer, Waltham, MA). Lipid accumulation was calculated as Nile Red RFU normalized to Hoechst RFU.

#### aP2 Induction

RNA was isolated using PureLink RNA Mini Kit (Thermo Scientific, Waltham, MA) and its concentration and purity were assessed in a NanoVue Spectrophotometer. Quantitative real time PCR was conducted using Power SYBR green RNA-to-Ct One-Step kit (Thermo Scientific, Waltham, MA) to assess the expression of adipogenesis-related genes.

#### PPAR-Responsive Element Prediction

A PPARgene database (ppargene.org) was used to predict the sequence specific PPAR-responsive elements (PPRE) on all the target genes measured in this study ^15^. Upon submitting the query, if the gene was predicted as a PPAR target gene, the query returned p-value and confidence level of the prediction and listed putative PPREs in the 5 kb transcription start site flanking region. Genes were assigned high-confidence category (*p* > 0.8), medium-confidence category (0.8 ≥ *p* > 0.6), and low-confidence category (0.6 ≥ *p* > 0.45). Genes with *p* value ≤ 0.45 were predicted as negative.

---

**Figure S1.**
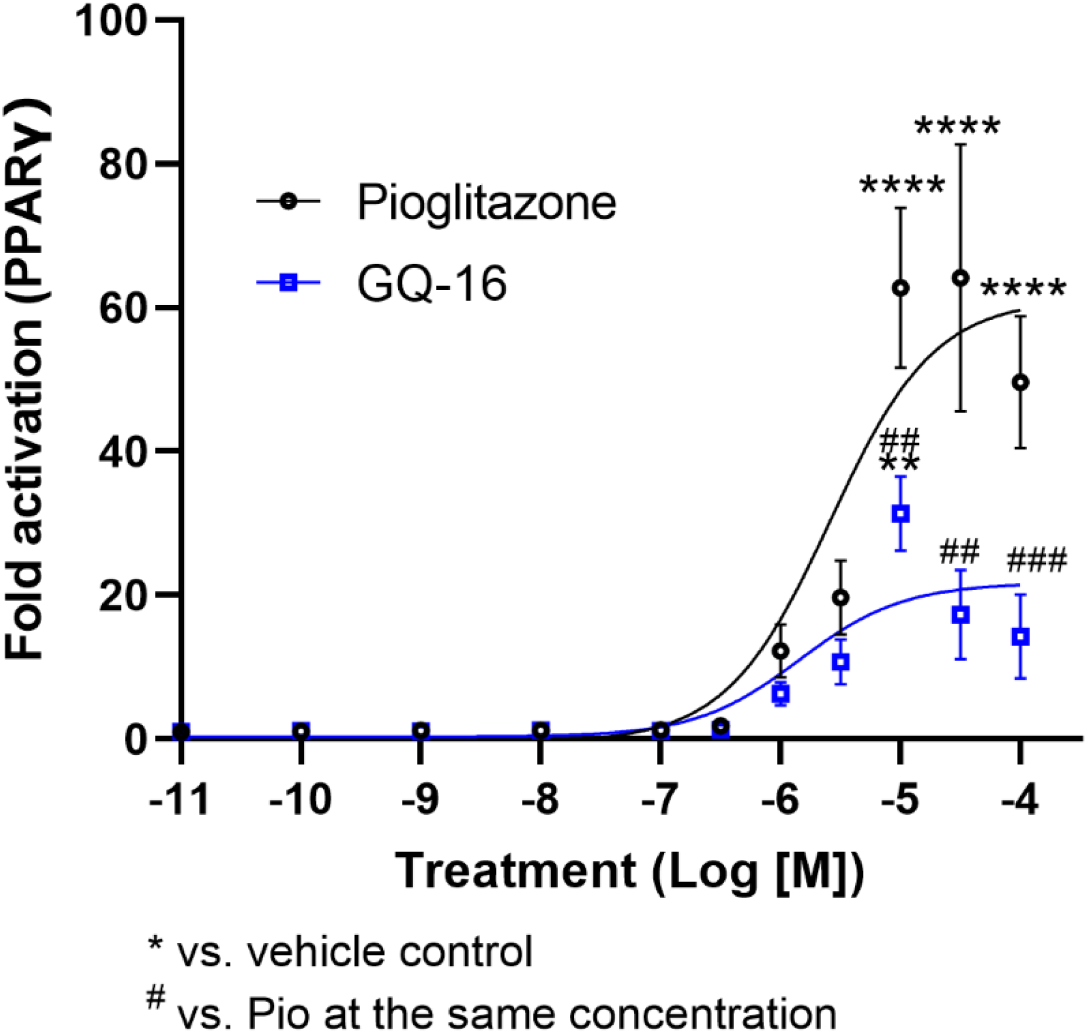
Dose Response Curve of PPARγ Activation with Pioglitazone and GQ-16. HeLa cells were transiently co-transfected with plasmids containing human *PPARG* Ligand-Binding Domain fused to the *GAL4* DNA-Binding Domain and a plasmid containing the luciferase reporter gene under the regulation of five Gal4 DNA-binding elements, treated with varying concentrations of Pio or GQ-16, and assayed for luciferase activity. While, GQ-16 and Pio activated PPARγ in a dose-dependent manner with EC50 1.4 vs 2.6 μM, respectively, GQ-16 elicited only ∼ 30-50% of the maximal activity induced by the full agonist Pio. * vs. vehicle control, ^#^ vs. Pio at the same concentration.

**Figure S2.**
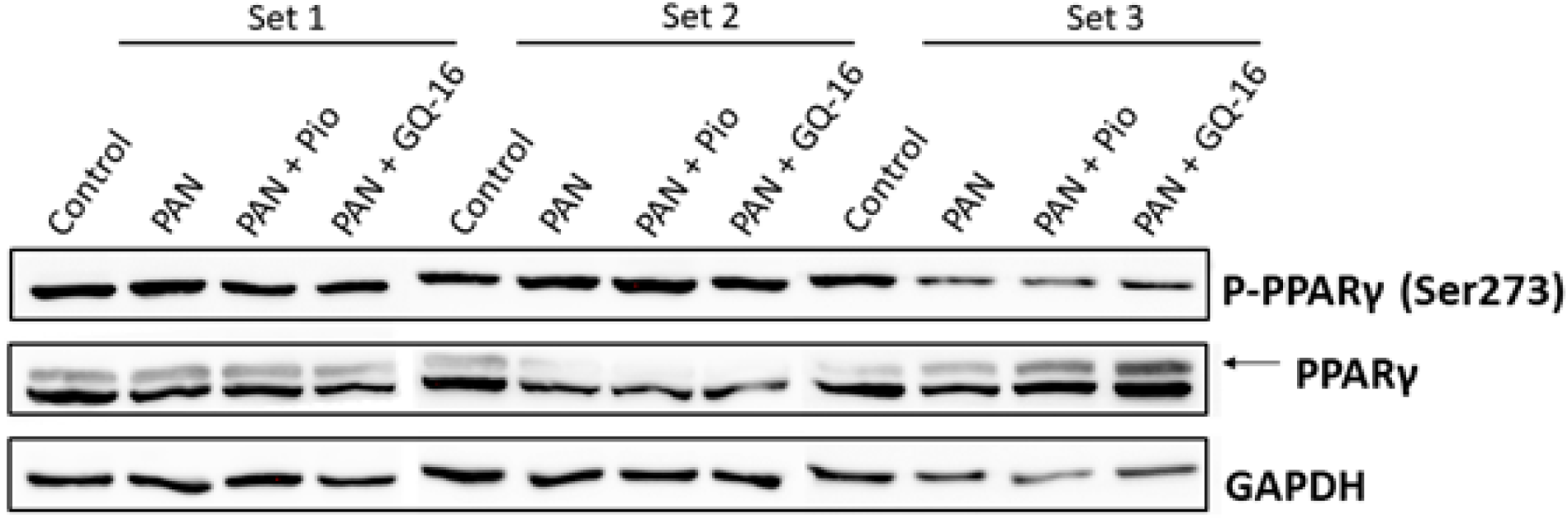
Phosphorylation Status of Glomerular PPARγ at Serine 273 Position. Total protein was isolated from Control, PAN-injected, and PAN-injected rats treated with Pio and GQ-16 from glomeruli. Western blotting was performed on the samples for P-PPARγ (Ser273), total PPARγ and GAPDH.

**Figure S3.**
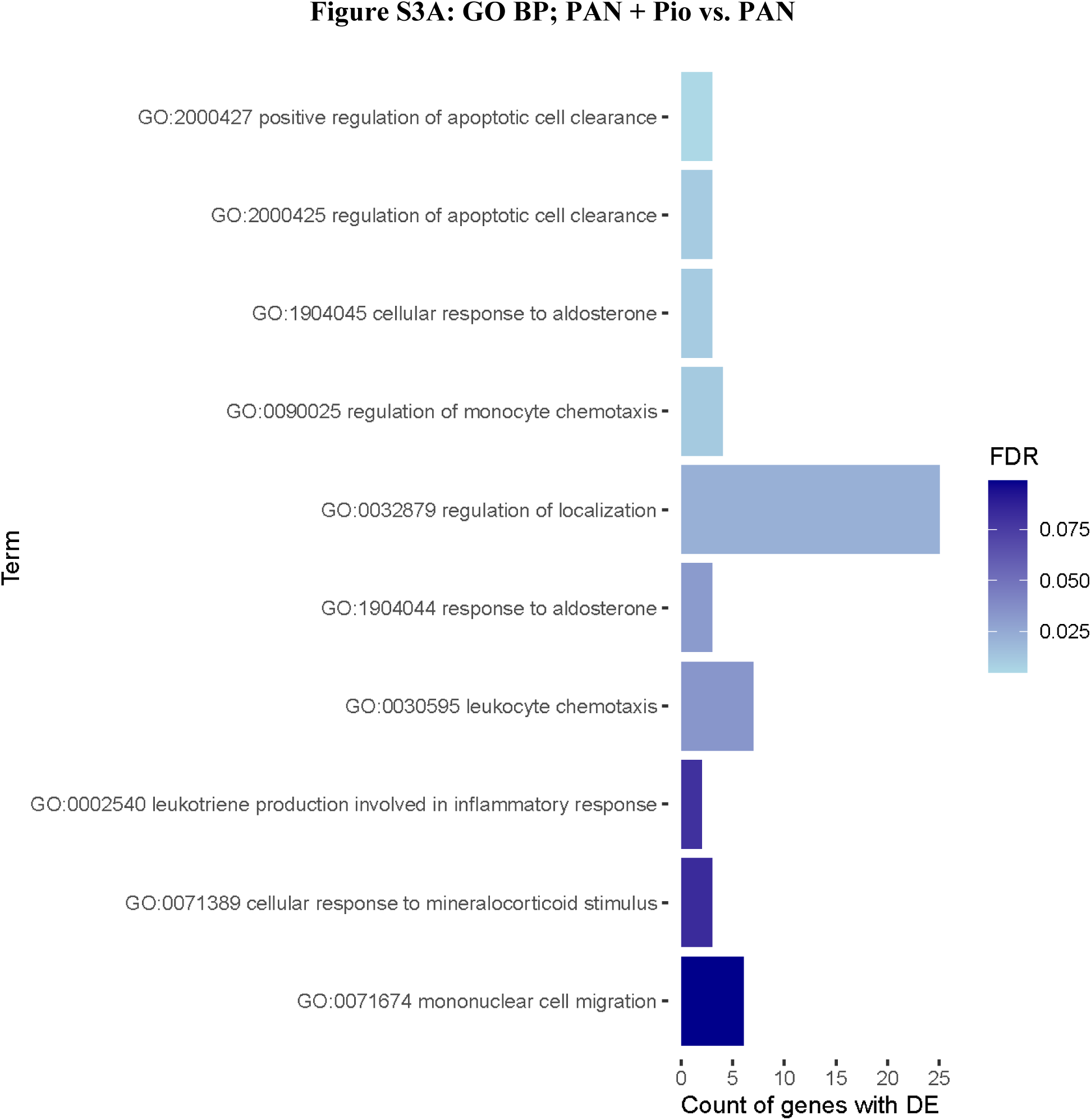
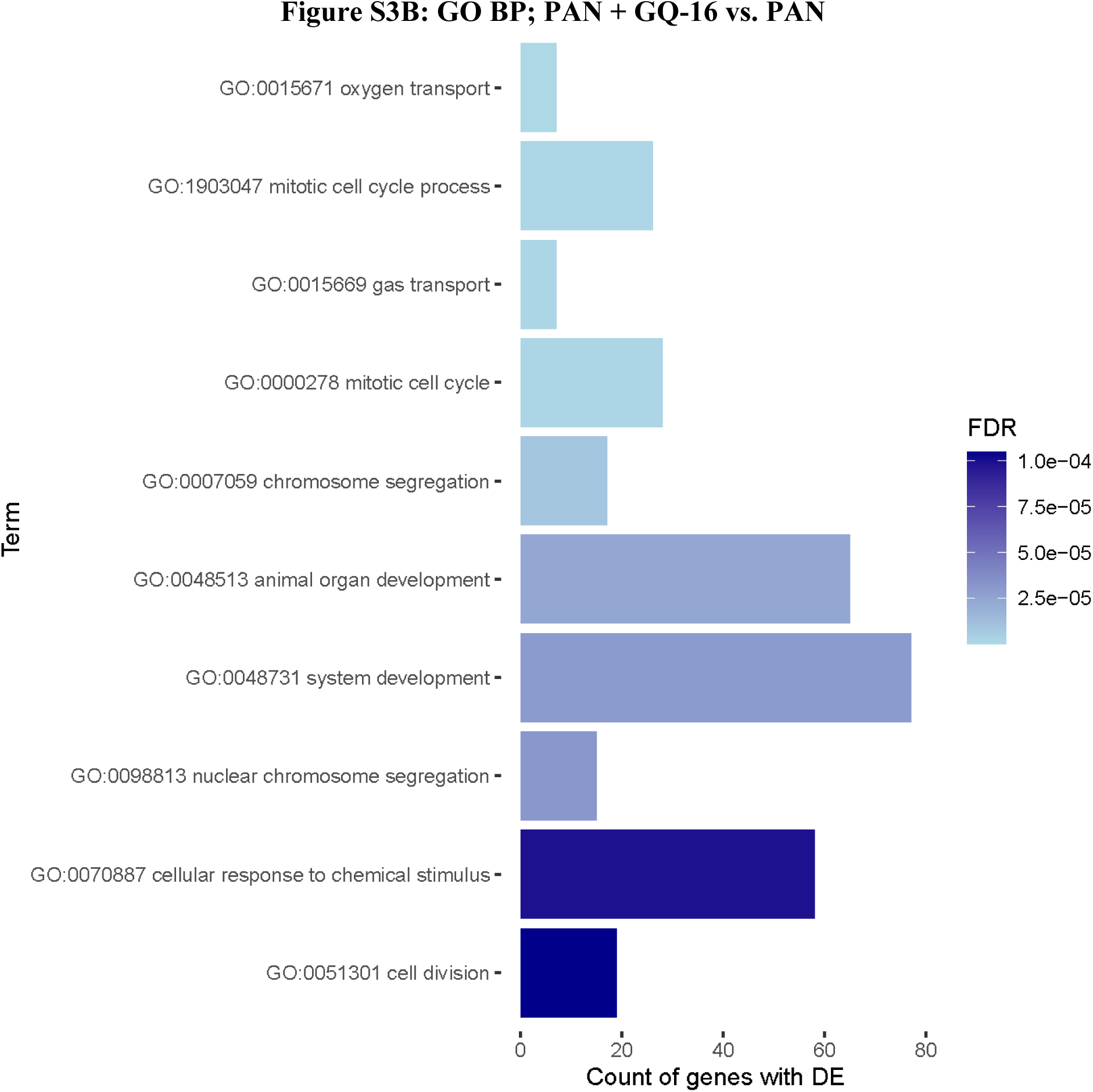
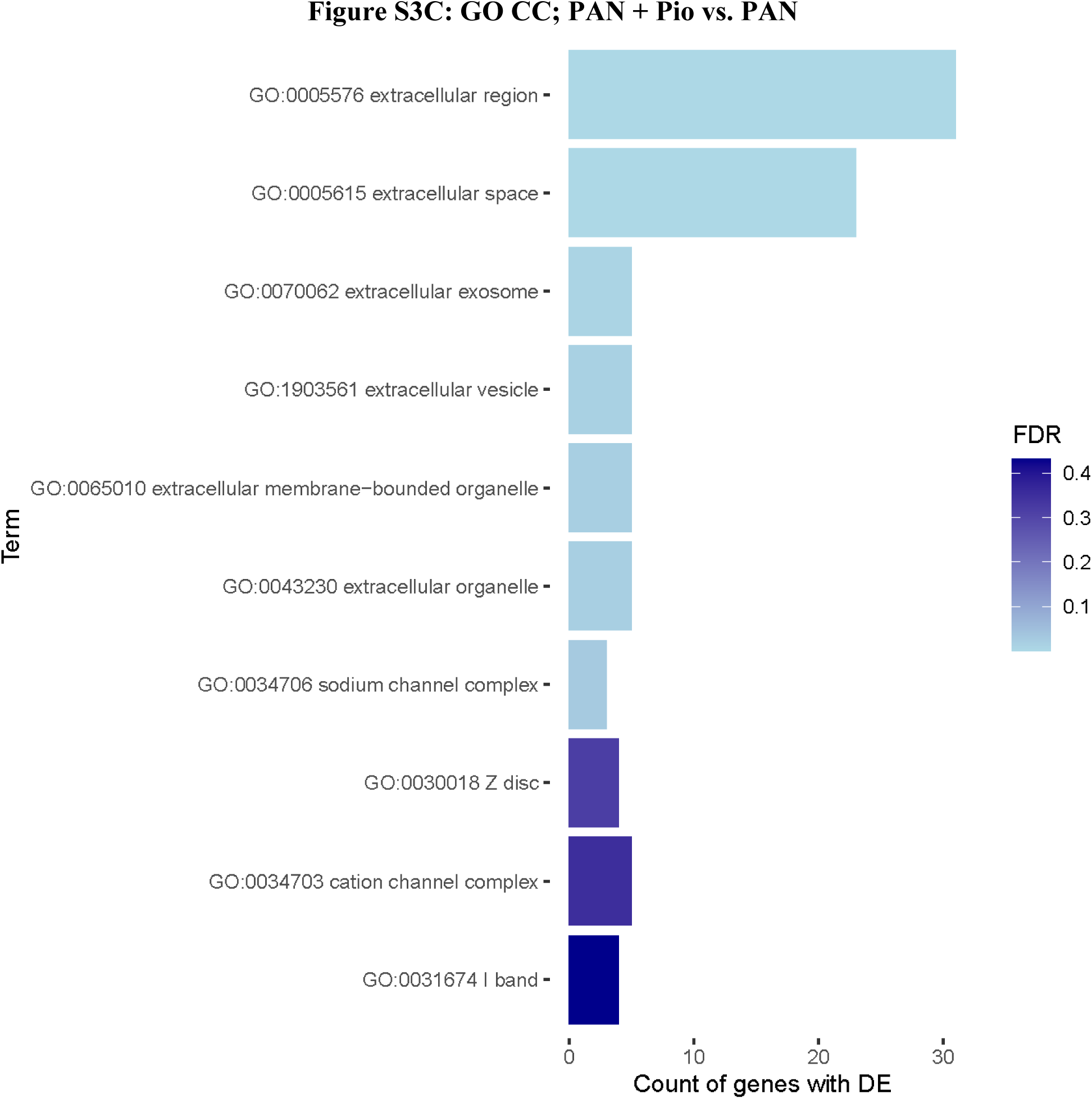
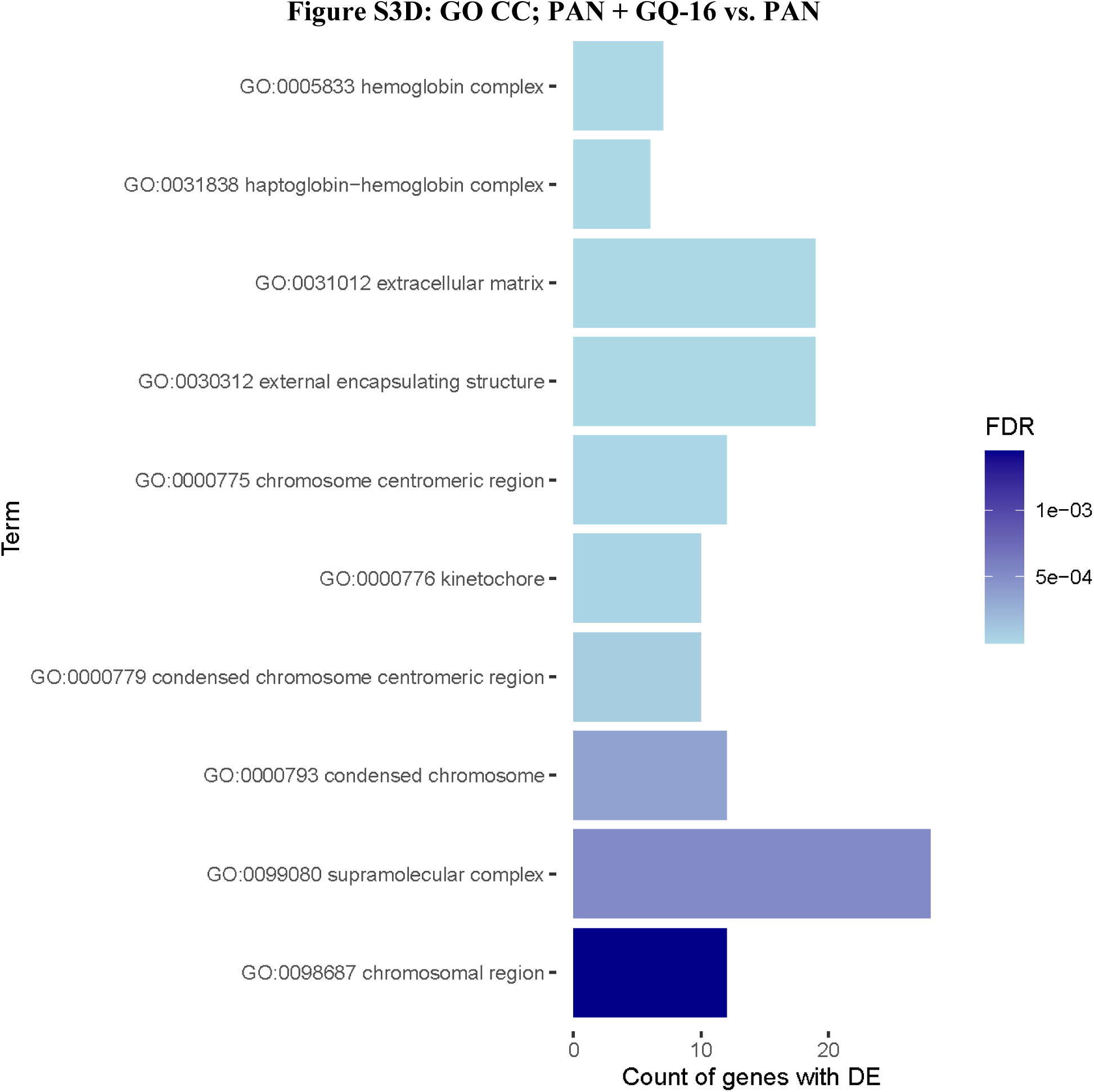
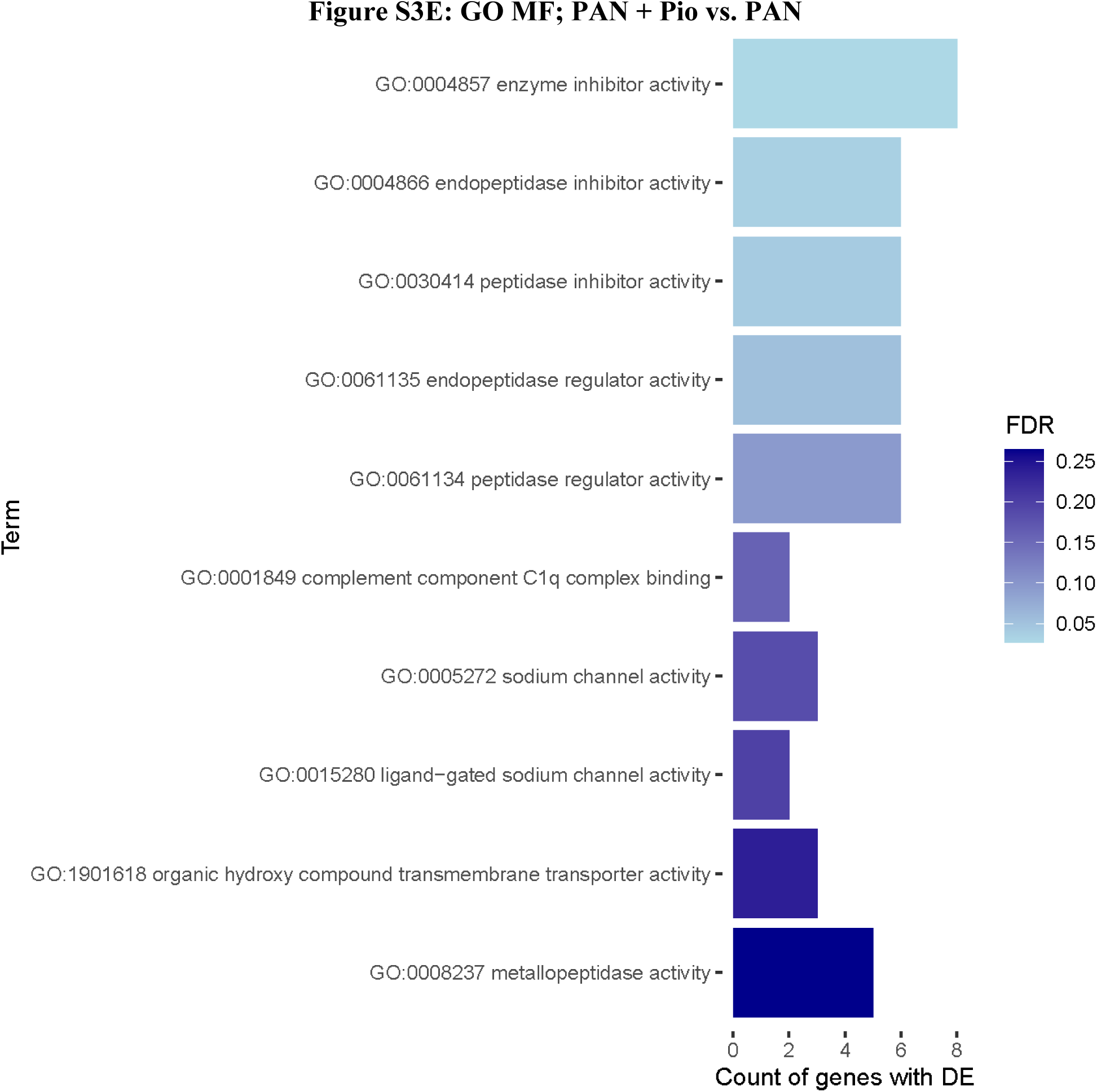
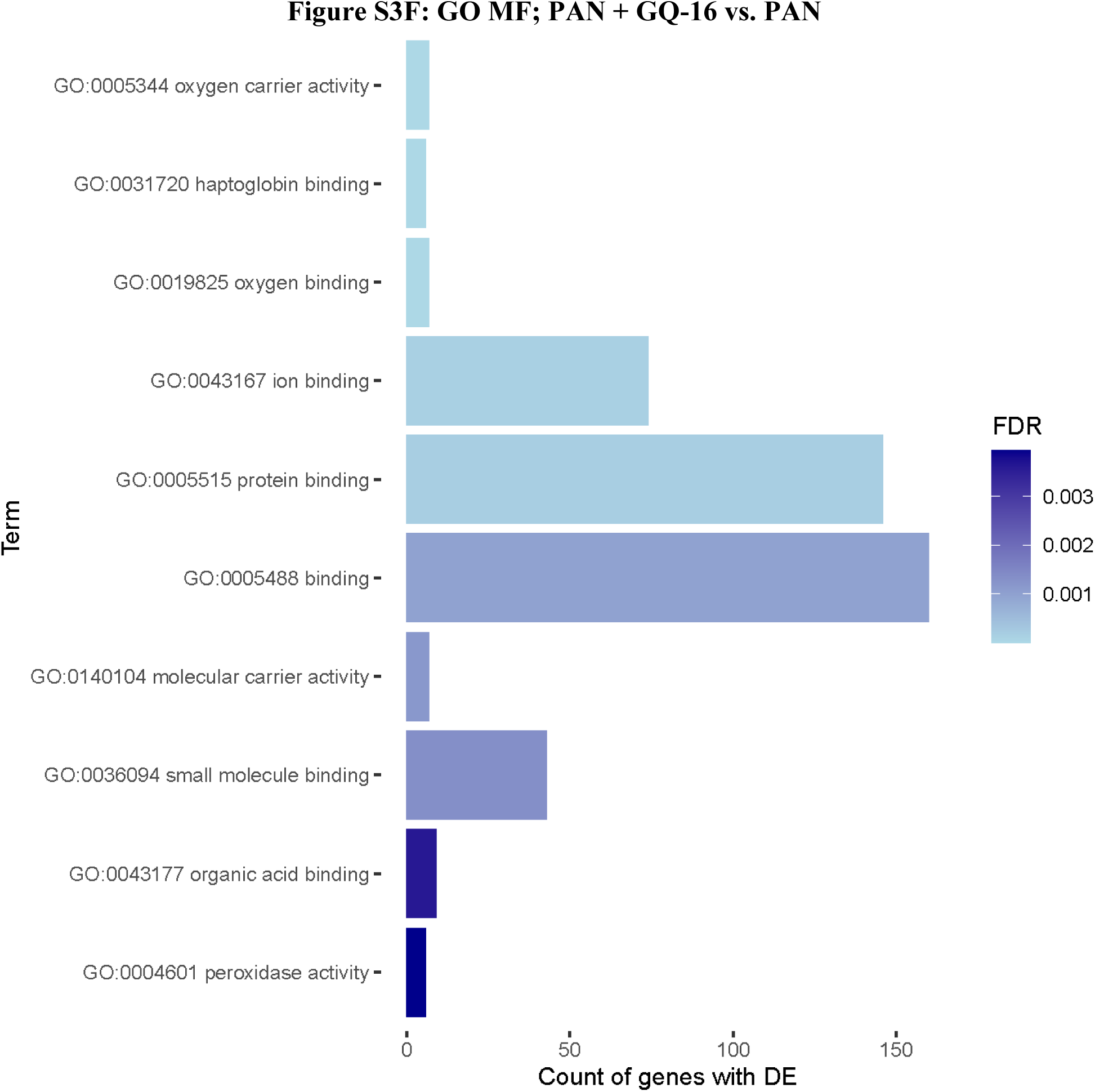
Ontology Enrichment Analysis. Ontology enrichment of the top 10 enriched GO terms for each comparison (PAN + GQ-16 vs. PAN and PAN + Pio vs. PAN) and for each GO type (Biological Processes [BP], cellular components [CC] and molecular functions [MF]) are plotted. **(A)** BP; PAN + Pio vs. PAN (4.27E-05 ≥ p ≥ 2.11E-06), **(B)** BP; PAN + GQ-16 vs. PAN (2.95E-08 ≥ p ≥ 1.28E-11), **(C)** CC; PAN + Pio vs. PAN (1.86E-03 ≥ p ≥ 8.14E-11) and **(D)** CC; PAN + GQ-16 vs. PAN (3.71E-06 ≥ p ≥ 1.82E-12), **(E)** MF; PAN + Pio vs. PAN (7.28E-04 ≥ p ≥ 7.39E-05), and **(F)** MF; PAN + GQ-16 vs. PAN (6.71E-06 ≥ p ≥ 1.41E-11). Color is based on FDR, length of bar is based on number of DEGs belonging to that GO term.

**Figure S4.**
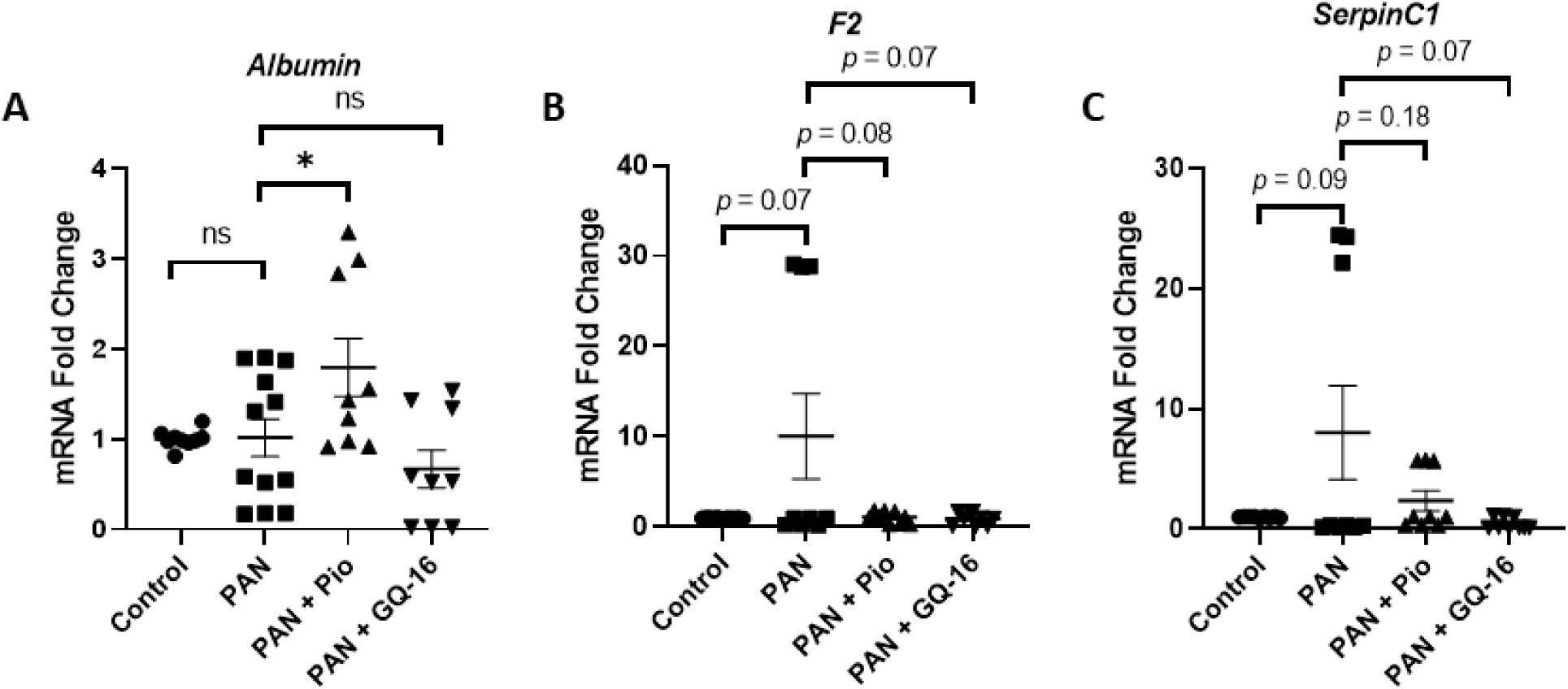
Hepatic Gene Expression of Albumin and Coagulation Factors F2 and SerpinC1. Real Time RT-PCR was used to measure gene expression from RNA isolated from liver tissue from Control, PAN-injected, and PAN-injected plus Pio or GQ-16 treatment rats. Mean ± SEM of **(A) *A****lbumin,* **(B)** *F2,* and **(C)** *SerpinC1* (n ≥ 3/group, assay in triplicates).

**Figure S5.**
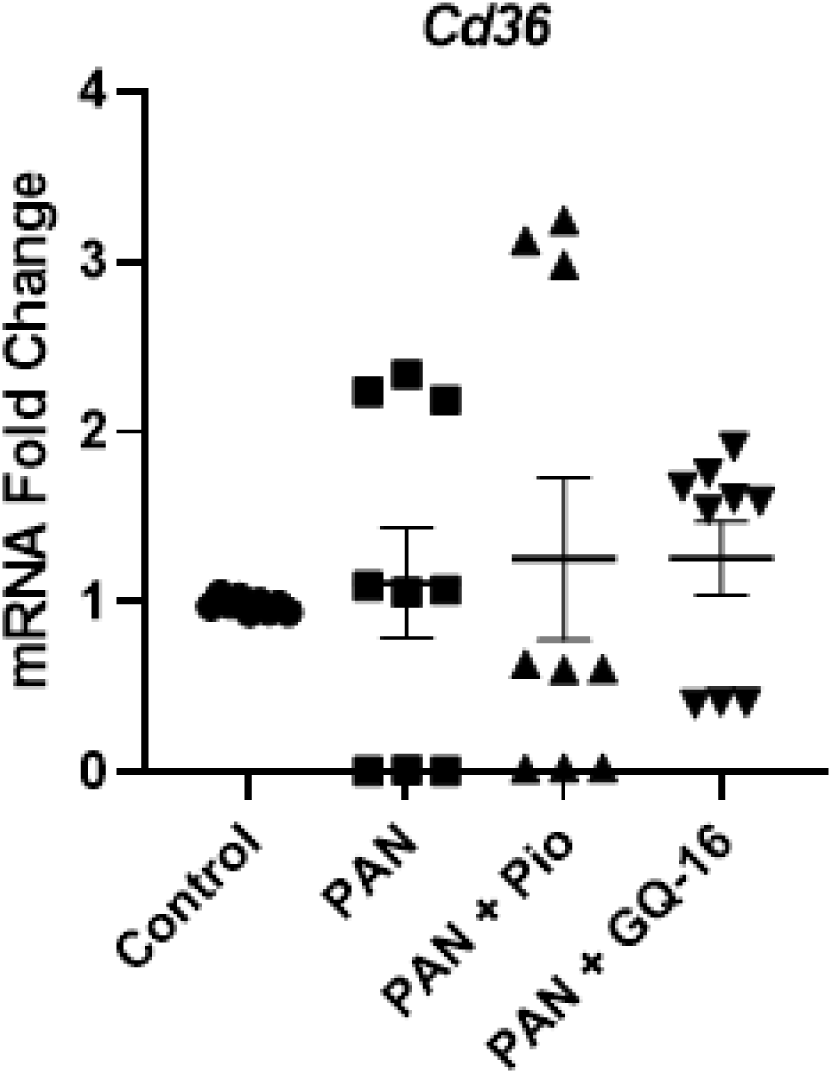
*WAT Cd36* Gene Expression. White adipose tissue gene expression was measured by real time RT-PCR from total RNA extracted from epididymal fat tissue from Control, PAN-injected, and PAN-injected rats treated with Pio and GQ-16 (n ≥ 3/group, assay in triplicates). Expression of mRNA from *Cd36* was determined and normalized to the fat house-keeping gene *Ppia*. Mean ± SEM plotted.

**Figure S6.**
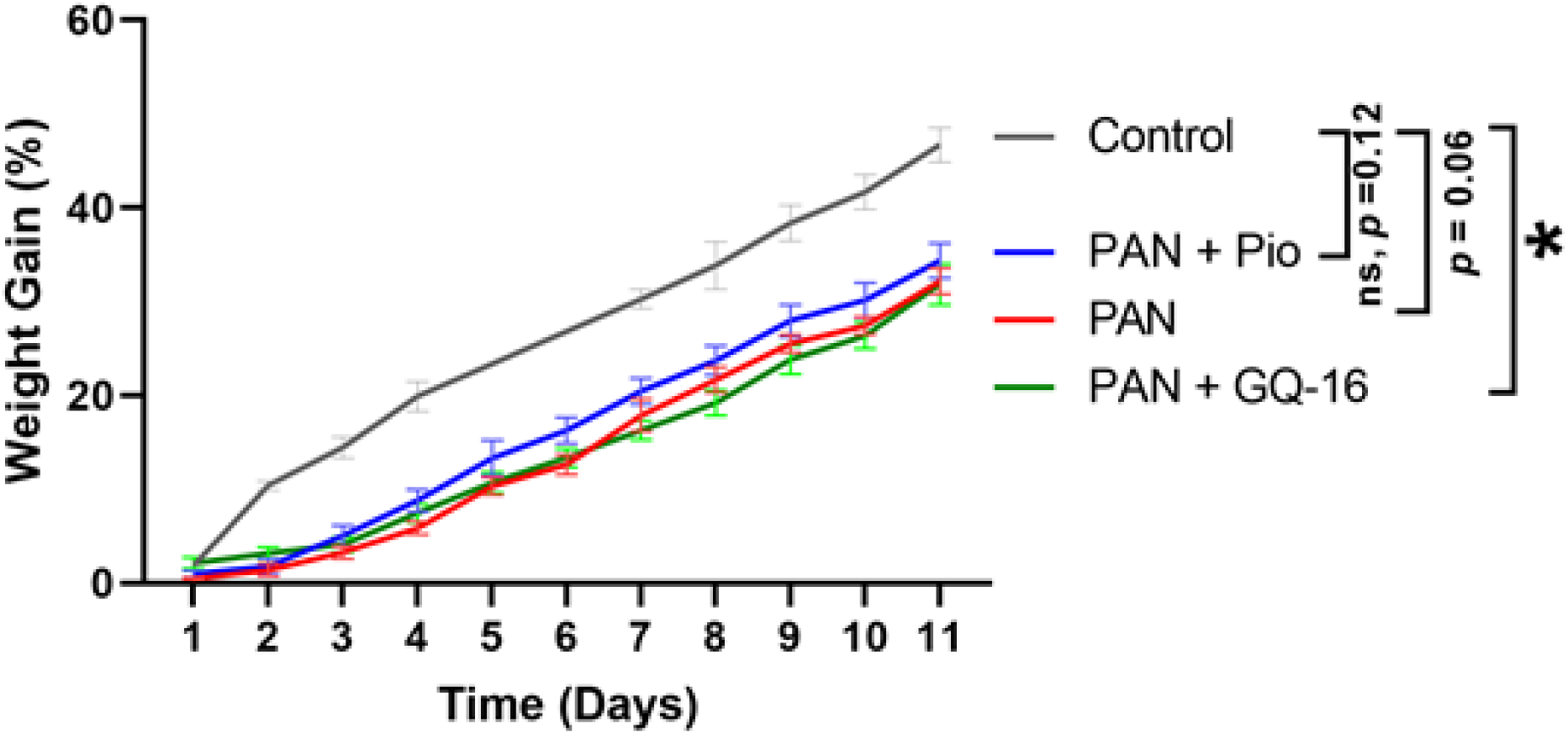
Percentage Weight Gain. Weight gain (%) was plotted with time from Control, PAN-injected and PAN-injected rats treated with Pio and GQ-16 (n=7/group).

